# Wnt target enhancer regulation by a CDX/TCF transcription factor collective and a novel DNA motif

**DOI:** 10.1101/2021.01.15.426889

**Authors:** Aravinda-Bharathi Ramakrishnan, Lisheng Chen, Peter Burby, Ken M. Cadigan

## Abstract

Transcriptional regulation by Wnt signalling is primarily thought to be accomplished by a complex of β-catenin and TCF family transcription factors (TFs). Although numerous studies have suggested that additional TFs play roles in regulating Wnt target genes, their mechanisms of action have not been investigated in detail. We characterised a Wnt-responsive element (WRE) downstream of the Wnt target gene *Axin2* and found that TCFs and Caudal-related homeodomain (CDX) proteins were required for its activation. Using a new separation-of-function TCF mutant, we found that WRE activity requires the formation of a TCF/CDX complex. Our systematic mutagenesis of this enhancer identified other sequences essential for activation by Wnt signalling, including several copies of a novel CAG DNA motif. Computational and experimental evidence indicates that the TCF/CDX/CAG mode of regulation is prevalent in multiple WREs. Put together, our results demonstrate the complex nature of cis- and trans- interactions required for signal-dependent enhancer activity.

## Introduction

The Wnt/β-catenin signalling pathway is highly conserved across the animal kingdom, plays numerous essential roles in animal development, and is required for the homeostasis of many tissues in adult organisms (Clevers and Nusse, 2012; Clevers et al., 2014). While it is well known that the Wnt pathway affects cell behaviour by transcriptionally regulating gene expression, many questions remain about how Wnt signalling controls gene expression in a cell-specific manner (Archbold et al., 2012; Lien and Fuchs, 2014; Ramakrishnan and Cadigan, 2017; Söderholm and Cantù, 2021).

The prevalent model of transcriptional regulation by the Wnt pathway is centred upon the regulation of β-catenin protein levels and the activity of transcription factors (TFs) of the TCF/LEF family (TCFs). In the absence of Wnt signals, β-catenin protein levels are kept low in the cells by a ‘destruction complex’, which targets β-catenin for proteasomal degradation. The binding of Wnt ligands to their extracellular receptor complexes inactivates the destruction complex, allowing β-catenin to accumulate (Stamos and Weis, 2013). β-catenin then complexes with TCFs, which are bound to specific TCF binding sites in cis-regulatory regions named Wnt-responsive elements (WREs) (Barolo, 2006). Subsequently, β-catenin recruits co-activators to regulate the transcription of Wnt target genes (Cadigan, 2012; Valenta et al., 2012).

The view of TCFs as the major transcriptional effectors of Wnt signalling is supported by genetic studies in vertebrates and invertebrates (van Es et al., 2012; Schweizer et al., 2003). More recently, studies in *Drosophila* and mammalian cell culture systems have confirmed that the majority of Wnt target genes fail to be activated in the absence of TCFs (Doumpas et al., 2018; Franz et al., 2017; Moreira et al., 2017). Functional TCF binding sites have been identified in many WREs, and synthetic reporters containing multimers of TCF sites are specifically activated by Wnt/β-catenin signalling (Archbold et al., 2012; Barolo, 2006). In light of this, TCFs are often considered necessary and sufficient for WRE activity. In addition to the High Mobility Group (HMG) domain that all TCFs possess, which binds to TCF binding sites (Cadigan and Waterman, 2012), invertebrate TCFs and some vertebrate TCF isoforms also contain an additional DNA-binding domain, the C-clamp (Atcha et al., 2007; Ravindranath and Cadigan, 2014). C-clamps recognizes GC-rich motifs termed Helper sites (Chang et al., 2008; Hoverter et al., 2014), which are essential for activation of multiple Wnt targets in Drosophila, C. elegans and mammalian systems (Archbold et al., 2014; Bhambhani et al., 2014; Chang et al., 2008; Hoverter et al., 2012).

The idea that WRE activity is solely mediated by TCFs is inconsistent with broader studies of enhancers, which suggest that these *cis-*regulatory elements are primarily regulated by combinatorial TF activity. Enhancers tend to contain clusters of binding sites for multiple TFs that vary in their number and relative orientation (Arnosti and Kulkarni, 2005; Lambert et al., 2018). In this manner, several TFs can be recruited to an enhancer through their cognate binding sites. There is evidence of protein-protein interactions between many TFs known to co-regulate enhancers. This has led to the idea of enhancer regulation by TF collectives, groups of TFs formed by protein-protein and protein-DNA interactions that act together to regulate transcription (Spitz and Furlong, 2012). Although theoretical frameworks suggest that a model of gene regulation involving TFs working in distinct collectives to activate enhancers can explain the observed specificity of gene regulatory networks, there are few examples of studies which have directly identified and tested the role of TF-TF interactions on gene expression (Lambert et al., 2018).

Consistent with the aforementioned view of enhancer structure, there are several lines of evidence that WREs are more complex than previously thought. While synthetic WREs consisting of high affinity TCF binding sites upstream of a minimal promoter (e.g., TOPflash) provide sensitive readouts for the Wnt pathway in cell culture, they do not faithfully recapitulate patterns of Wnt signalling at the organismal level (Barolo, 2006; Chang et al., 2008). Reporters knocked into endogenous Wnt target genes such as Axin2 have proven to be better markers for Wnt signalling (van Amerongen et al., 2012), suggesting that the endogenous WREs regulating them contain additional information and are not just collections of TCF binding sites. Similarly, a recent study in *Xenopus* found that β-catenin recruitment to chromatin was insufficient for the activation of nearby genes, leading the authors to suggest that additional TFs are required at WREs to act as spatio-temporal specificity cues (Nakamura and Hoppler, 2017; Nakamura et al., 2016).

Correlative experiments using chromatin immunoprecipitation (ChIP) based techniques have identified several TFs that show significant levels of chromatin co-occupancy with TCFs (Bhambhani and Cadigan, 2014; Söderholm and Cantù, 2021). Some of these TFs also directly interact with TCFs, suggesting that they could be part of a Wnt/TCF TF collective. For instance, TCF7L2 and CDX2 show significant levels of chromatin co-occupancy in colorectal cancer cells (Verzi et al., 2010), and protein-protein interactions have been observed between other members of the TCF and CDX families (Béland et al., 2004a). Similarly, the TFs Sp5 and Sp8 interact with multiple TCFs and bind to several WREs in mouse embryonic stem cells (Kennedy et al., 2016). Co-occupancy at genomic locations and interactions between TCFs and other TFs have also been reported, including Smads and AP-1 (Archbold et al., 2012), as well as TEADs (Jiao et al., 2017). Together, these studies suggest the existence of multiple TCF-containing TF collectives which contribute to the high cell-type specificity seen in Wnt target genes. However, the importance of interactions between TCFs and other TFs has not been rigorously tested, and whether there is a TF “binding site grammar” shared by WREs has not been systematically examined.

In this study, we have analysed a novel WRE located downstream of the *Axin2* Wnt target gene in great detail. We found that in addition to TCF proteins, enhancer activity was regulated by Caudal-related homeodomain (CDX) protein levels and that TCF7L2 and CDX1 are recruited to this WRE. We found evidence for the existence of a TCF7-CDX1 complex, and our experiments using a specific separation-of-function mutant of TCF7 support a model where TCF7-CDX1 complex formation is required for enhancer activity. Systematic scanning mutagenesis of the WRE revealed that as is typical for WREs, it contains four TCF binding sites that are absolutely required for activation. In addition, it identified other sequences required for WRE activity, including two sequence motifs resembling the CDX consensus sequence and a previously unidentified orphan DNA motif we term as CAG sites. Functional TCF, CDX, and CAG sites are also present in a WRE linked to human cancers located upstream of the *c-Myc* oncogene. Computational analysis of chromatin bound by TCF7L2 and CDX2 revealed an enrichment of CAG motifs, suggesting that a TCF/CDX/CAG cassette exists in other WREs. Using synthetic reporters, we also saw that the CDX and CAG sites do not respond to Wnt/β-catenin signalling, but these motifs can potentiate the ability of TCF sites to respond to pathway activation in a cell-type specific manner. These results define a binding site grammar for a subset of WREs, and provide a more detailed view of their structure beyond the simplistic model of WREs as clusters of TCF sites.

## Results

### Identification and characterisation of CREAX, a highly Wnt-regulated enhancer near *Axin2*

To test the features of the TF collective model in the context of Wnt signalling, we looked for highly responsive WREs which could be studied under cell culture conditions. *Axin2* is a robustly activated Wnt target gene in several mammalian tissues (Lim et al., 2016; van Amerongen et al., 2012; Wang et al., 2015) and cell lines (Jho et al., 2002; Leung et al., 2002; Van der Flier et al., 2007). A recent β-catenin ChIP-seq study found two TCF-dependent β-catenin-bound regions near the *Axin2* gene locus (Doumpas et al., 2018), one near the promoter and one located about 45 kb downstream of the TSS (Fig. 1A). Previous studies found peaks of TCF7L2 binding at both loci (Mokry et al., 2010). The promoter-proximal region, termed Ax2, was found to be responsive to Wnt signalling in a reporter assay in HEK293T cells (Jho et al., 2002). The distal enhancer, which we named CREAX (Cdx Regulated Enhancer near Axin2), was shown to be active in the LS174T colorectal cancer cell line (Hatzis et al., 2008). We cloned both elements into pGL4.23 vectors upstream of a minimal TATA-box promoter and luciferase (luc) gene and examined their activity in HEK293T cells with and without Wnt signalling.

**Fig 1:**
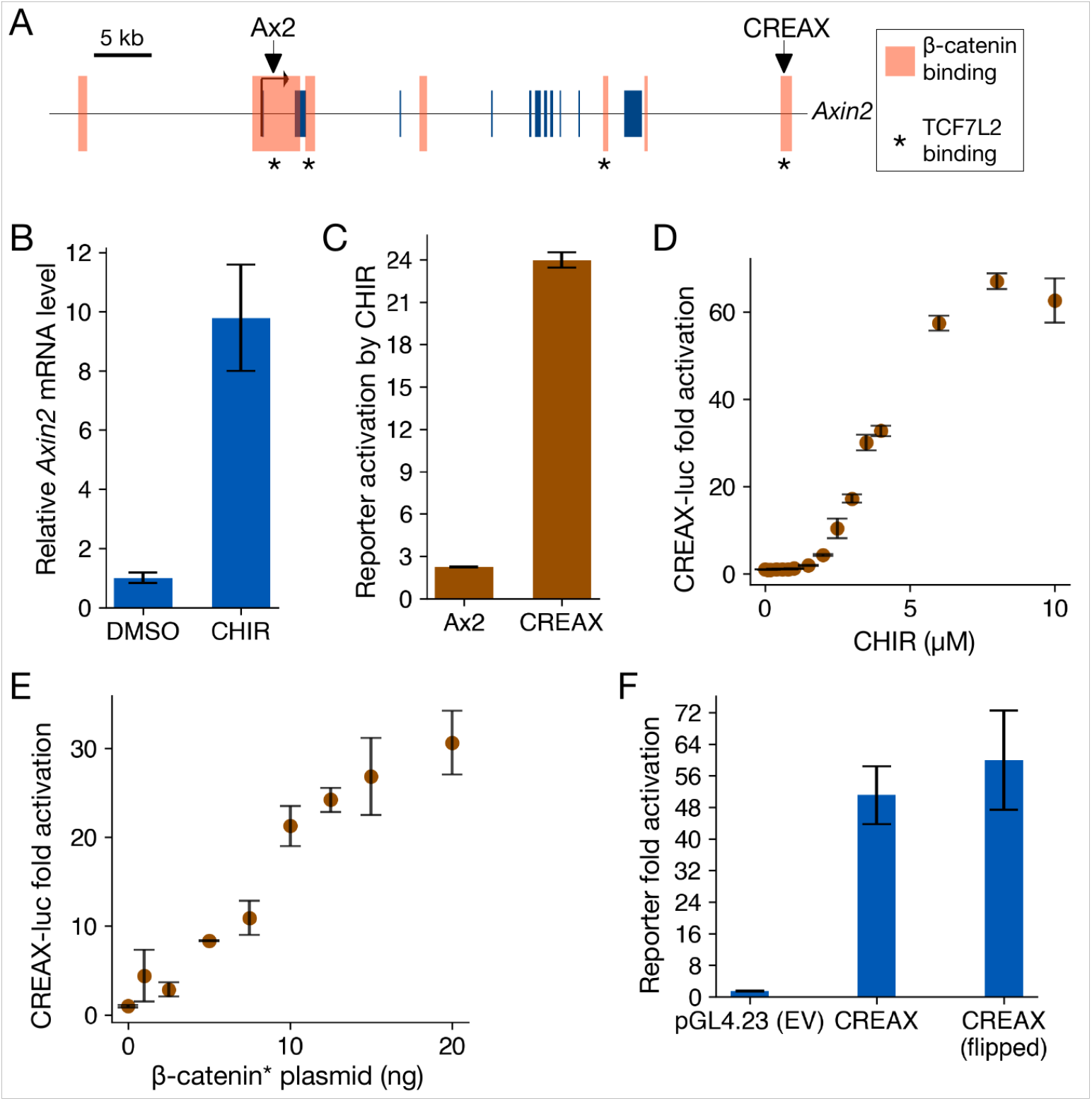
Characterisation of CREAX (Cdx Regulated Enhancer near Axin 2), a highly Wnt-responsive enhancer near *Axin2.* A) Position of CREAX (Cdx Regulated Enhancer near Axin2) relative to the Axin2 gene locus. Smaller blue boxes represent exons and larger orange boxes are regions bound by β-catenin in LS174T cells (Mokry et al., 2010). Known TCF7L2-bound regions are denoted by asterisks (Doumpas et al 2018). Positions are shown with respect to the hg18 assembly. B) RT-qPCR measurement of Axin2 transcript levels showing an increase upon treatment with CHIR-99021 (CHIR) in HEK293T cells. C) Activity levels of luciferase reporters of the previously identified Ax2 promoter-proximal WRE (Jho et al., 2002) and CREAX with CHIR treatment shows that CREAX is more sensitive to Wnt signalling. D) CREAX-luciferase reporter activity increases in a dose-dependent manner to CHIR concentration in HEK293T cells. E) CREAX-luciferase reporter activity increases in a dose-dependent manner upon transfection of increasing amounts of a plasmid expressing stabilised β-catenin containing the S33Y mutation (β-catenin*). F) Activation of the CREAX-luciferase reporter by β-catenin* is not affected by orientation, meeting the classical definition of an enhancer. Data information: In (B-F), data are presented as mean ± SD from three replicates (N = 3) for each condition.

Activating the Wnt/β-catenin pathway in HEK293T cells using the GSK3 inhibitor CHIR-99021 (CHIR) resulted in a 10-fold increase in *Axin2* transcript levels as detected by RT-qPCR (Fig. 1B). The same treatment caused a modest increase in activity of the Ax2-luciferase reporter but a much stronger increase in the CREAX-luciferase reporter (Fig. 1C). CREAX-luc activity increased in a dose-dependent manner with increasing concentrations of CHIR (Fig. 1D). Similarly, CREAX-luc showed dose-responsive activation when co-transfected with increasing amounts of a plasmid expressing β-catenin containing the stabilising S33Y mutation (β-catenin*, Fig. 1E) which prevents it from being targeted by the destruction complex (Stamos and Weis, 2013). Finally, we found that CREAX activity was not affected by reversing its orientation, meeting the standard requirement for an enhancer (Fig. 1F). The high amplitude of response to Wnt/β-catenin signalling provided a strong starting point for a detailed analysis of the TFs and *cis-*regulatory sequences that regulate the activity of this WRE.

### TCF and CDX family proteins bind to and regulate CREAX and a distal enhancer of *c-Myc*

To identify the TFs that regulate CREAX activity, we computationally scanned the sequence of the enhancer for putative binding sites of TFs known to co-localise with TCFs at WREs. We used motif information from the JASPAR database (Fornes et al., 2020) along with the FIMO utility from the MEME suite (Grant et al., 2011) to perform this search. This preliminary analysis identified potential binding sites for TCF and CDX family TFs in CREAX. We found no sequences resembling Helper sites, the DNA motif recognized by TCF7 and TCF7L2 isoforms containing the C-clamp DNA binding domain (Atcha et al., 2007; Hoverter et al., 2012, 2014). A similar scan of the Ax2 WRE sequence identified putative TCF binding sites, but none for CDX. To confirm a role for CDX proteins in regulating CREAX, we activated Wnt signalling in HEK293T cells with β-catenin* and examined the effect of RNAi against CDX and TCF TFs on CREAX-luc reporter activity. RNAi against CDX1 or the TCFs LEF1 and TCF7 reduced CREAX reporter activity (Fig. 2A).

**Fig 2:**
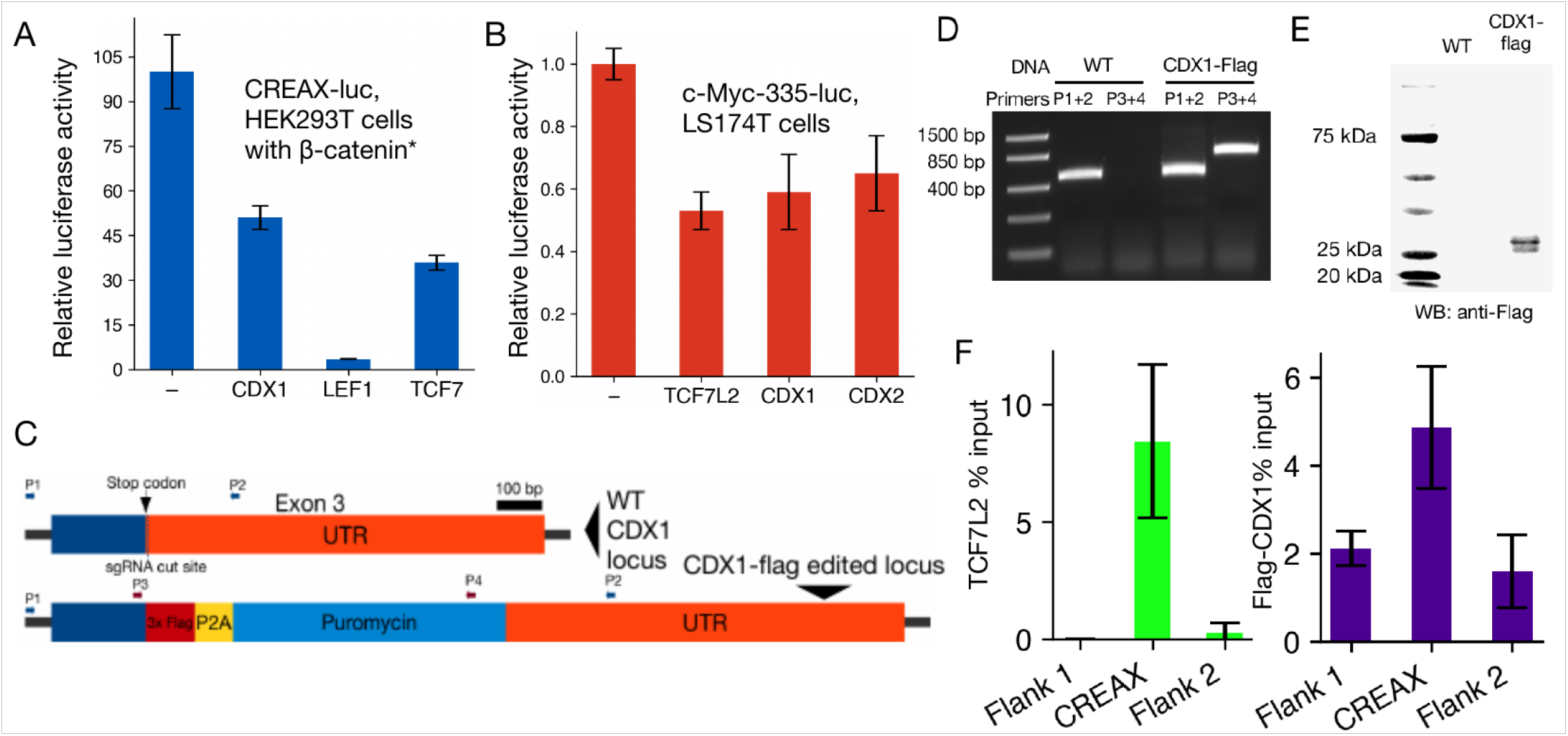
CDX and TCF family members directly bind to and regulate the activity of CREAX and a distal enhancer of c-Myc. A) RNAi-mediated depletion of CDX1, LEF1, or TCF7 reduces the activation of CREAX-luciferase by β-catenin* in HEK293T cells. Cells were transfected with CREAX-luc, β-catenin* and CDX1, LEF1, or TCF7 shRNA expression plasmids. B) Activity of a reporter of the c-Myc-335 enhancer previously shown to be bound by TCF7L2 and CDX2 in LS174T cells is reduced by RNAi against TCF7L2, CDX1, or CDX2. Cells were transfected with the reporter and plasmids expressing shRNA targeting TCF7L2, CDX1, or CDX2. C) Cartoon showing a portion of the CDX1 gene locus in HEK293T cells edited to express C-terminally Flag-tagged CDX1. The different sets of primers used to validate the edits are shown with arrows. D) PCR validation showing that the CDX1-flag cells are heterozygous for the CDX1-flag allele. E) Anti-Flag western blot showing the expression of Flag-tagged protein in the CDX1-flag cells. F) TCF7L2 and CDX1 both bind to CREAX. Anti-TCF7L2 and anti-Flag ChIP was performed on Flag-CDX1 HEK293T cells after treatment with CHIR. ChIP signals were measured using qPCR with primers targeting CREAX or flanking regions on both sides and normalised to signal from input chromatin. In (A,B,F), data are shown as mean ± SD from three replicates (N = 3) for each condition.

After examining previously studied WREs to find ones that might be co-regulated by TCF and CDX TFs, we identified c-Myc-335, a distal upstream enhancer of the *c-Myc* oncogene (Lewis et al., 2014; Verzi et al., 2010). This WRE had been of interest in cancer biology since it can contain a naturally occurring SNP (rs6983267) linked to increased colorectal and prostate cancer risk (Tomlinson et al., 2007; Zanke et al., 2007). This polymorphism was found to reside in a TCF binding site and increase its affinity to TCF7L2 (Pomerantz et al., 2009; Tuupanen et al., 2009; Wright et al., 2010). To test whether this WRE was regulated in a similar manner to CREAX, we constructed a c-Myc-335 luciferase reporter. We tested it in LS174T colorectal cancer cells, which have mutations in β-catenin that result in constitutively active Wnt signalling (van de Wetering et al., 2002). In these cancer cells, RNAi against TCF7L2, CDX1, and CDX2 reduces the activity of the c-Myc-335 reporter (Fig. 2B).

Most previous literature on the subject of TCF/CDX co-occupancy focuses on CDX2 in colorectal cancer cells (Lewis et al., 2014; Verzi et al., 2010). Since CREAX activity in HEK293T cells was sensitive to CDX1 protein levels, we were interested in whether CDX1 was also capable of co-occupying WREs along with TCFs. To this end we used ChIP-qPCR to test the binding of CDX1 to CREAX in HEK293T cells. Since we were unable to find ChIP-quality antibodies against CDX1, we generated a polyclonal HEK293T cell line expressing C-terminally flag-tagged CDX1 protein from the endogenous gene locus. To create this line, we adapted the previously reported CETCh-seq protocol (Savic et al., 2015) to generate a polyclonal HEK293T cell population using CRISPR/Cas9 mediated genome editing that express a Flag-tagged CDX1 protein. The edit replaced the endogenous stop codon with a cassette containing a 3xFlag epitope upstream of the coding sequences for the P2A self-cleaving peptide and the puromycin resistance gene (Fig. 2C). After selecting for puromycin resistance, we obtained a population of edited HEK293T cells which expressed a flag-tagged CDX1 detectable on a western blot (Fig. 2E). PCR using combinations of primers located inside and outside the Flag epitope showed that the polyclonal cell population contains a mixture of both the tagged and untagged alleles (Fig. 2C,D).

Using the CDX1-Flag cell line, we performed ChIP using antibodies against TCF7L2 and the Flag tag (for CDX1). We conducted the experiments after activating Wnt signalling using CHIR to examine chromatin state under conditions in which the enhancer was active. To detect the binding of TCF7L2 and CDX1, we used qPCR with primers located in the enhancer and compared the results to the signal from primers in flanking regions outside the enhancer. Both TCF7L2 and CDX1 were highly enriched at the CREAX locus, with the TCF7L2 ChIP signal being 30-fold higher from the enhancer compared to the flanks, and CDX1 showing a 2.3-fold higher signal at the enhancer (Fig. 2F).

### Direct TCF-CDX protein-protein interactions are required for enhancer function

Previous reports indicated that LEF1 and CDX1 could form a complex and showed that residues on the basic tail of LEF1 were required for this interaction (Béland et al., 2004b). The basic tail is a short stretch of basic amino acid residues found in all TCFs located immediately C-terminal to the HMG domain (Archbold et al., 2012) (Fig. 3A). These residues are conserved in other TCFs (Fig. 3B). To test the role of TCF-CDX interactions in enhancer activity, we attempted to generate TCF proteins with mutations preventing their interaction with CDX TFs. As the N-terminal portion of the basic tail is involved in contacting DNA (Giese et al., 1991), we focused on residues at the C-terminal end of the motif. We made charge-swap mutations in two residues (R350E, K352E) of a TCF7 isoform lacking a C-clamp to generate a variant we named BTmut (basic tail mutant). We transfected HEK293T cells with plasmids expressing Flag-tagged CDX1 and HA-tagged WT TCF7 or BTmut. We found that TCF7 robustly co-immunoprecipitated CDX1 and BTmut did not (Fig. 3C), suggesting that the mutations severely compromised the ability of BTmut to complex with CDX1. This result was also confirmed by a pulldown assay with recombinant GST-tagged CDX1 protein and His-tagged TCF7 or BTmut (Fig. 3D). These results suggested that BTmut could be a useful tool for testing the importance of TCF-CDX interactions in WRE activation.

**Fig 3:**
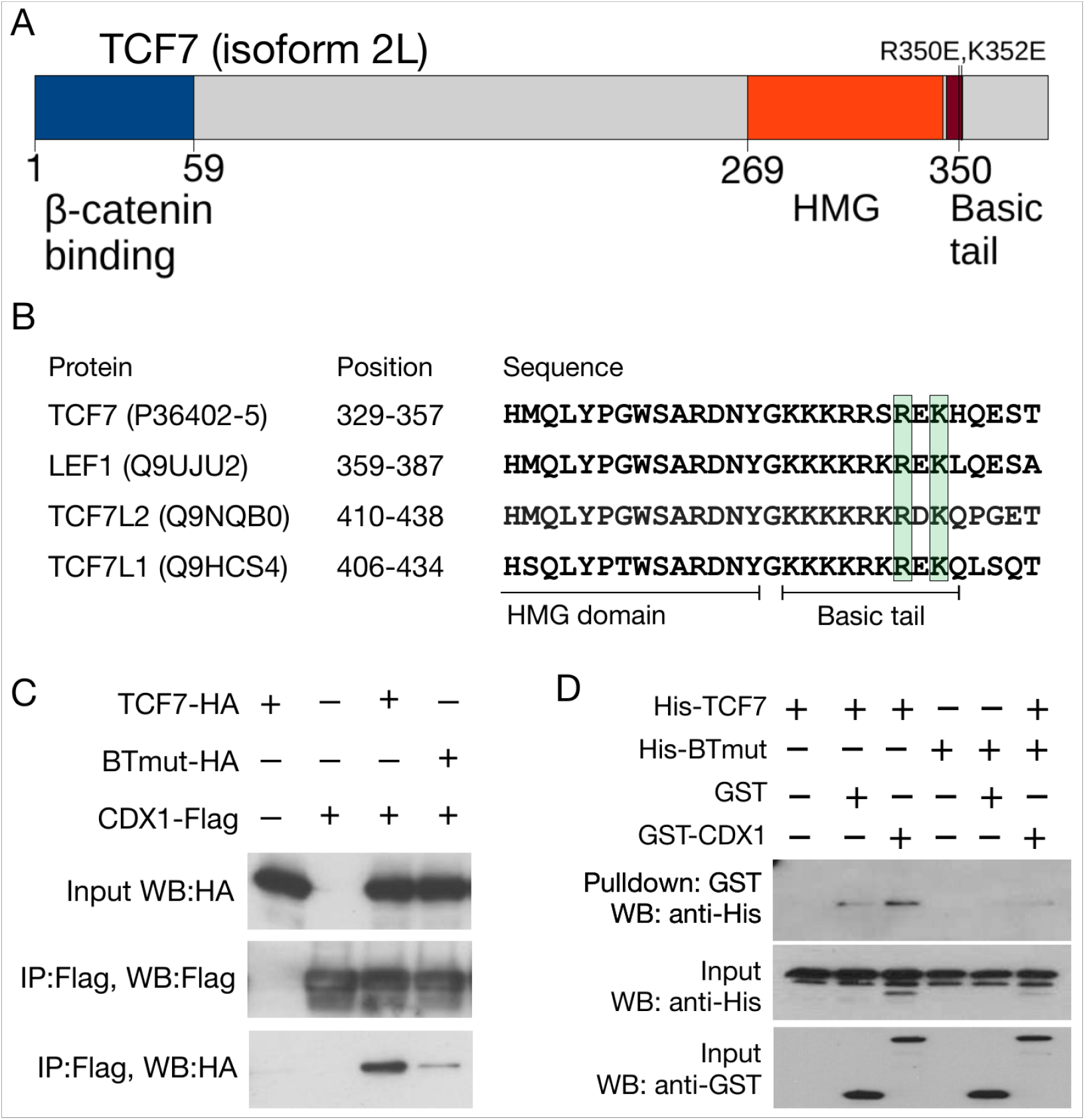
TCFs bind to CDX proteins through highly conserved residues in the basic tail. A) Cartoon of TCF7 protein with functional domains highlighted. Positions of R350E and K352E, mutations which interfere with the TCF-CDX interaction, are highlighted. B) Alignment of the C-terminal portion of the HMG domain and basic tail of the 4 human TCFs showing conservation of R350 and K352. Uniprot identifiers for the proteins are shown along with positions of the residues in the alignment. C) Co-IP showing that the BTmut variant of TCF7 with 2 amino acid substitutions (R350E,K352E) shows attenuated binding to CDX1. HEK293T cells were transfected with CDX1-flag and HA-tagged WT or BTmut variants of TCF7. Cell lysates were subjected to an anti-Flag IP and examined using anti-HA and anti-Flag western blots (WB). Input samples were used as loading control. D) GST pulldown showing direct binding between recombinant CDX1 and WT TCF7 but not BTmut. Purified His-tagged TCF7 or BTmut proteins were pulled down using glutathione beads with GST or GST-tagged CDX1 as bait. Precipitates were analysed on an anti-His WB.

In order for BTmut to be an informative reagent for probing the importance of TCF7-CDX1 binding in transcriptional regulation, we needed to ensure that the mutations in the basic tail did not interfere with other essential functions of TCF7. To this end, we compared the activity of WT and BTmut TCF7 proteins using the TOPflash reporter in HEK293T cells, a synthetic reporter containing 6 multimerised TCF sites (Korinek et al., 1997). RNAi against LEF1 reduced the activation of TOPflash by β-catenin*. Consistent with previous work in the field which suggests that the 4 mammalian TCFs bind to very similar DNA motifs and can be functionally redundant (Archbold et al., 2012; Moreira et al., 2017), we found that in the case of TOPflash, the LEF1 RNAi effect could be rescued by the overexpression of TCF7 (Fig. 4A).

**Fig 4:**
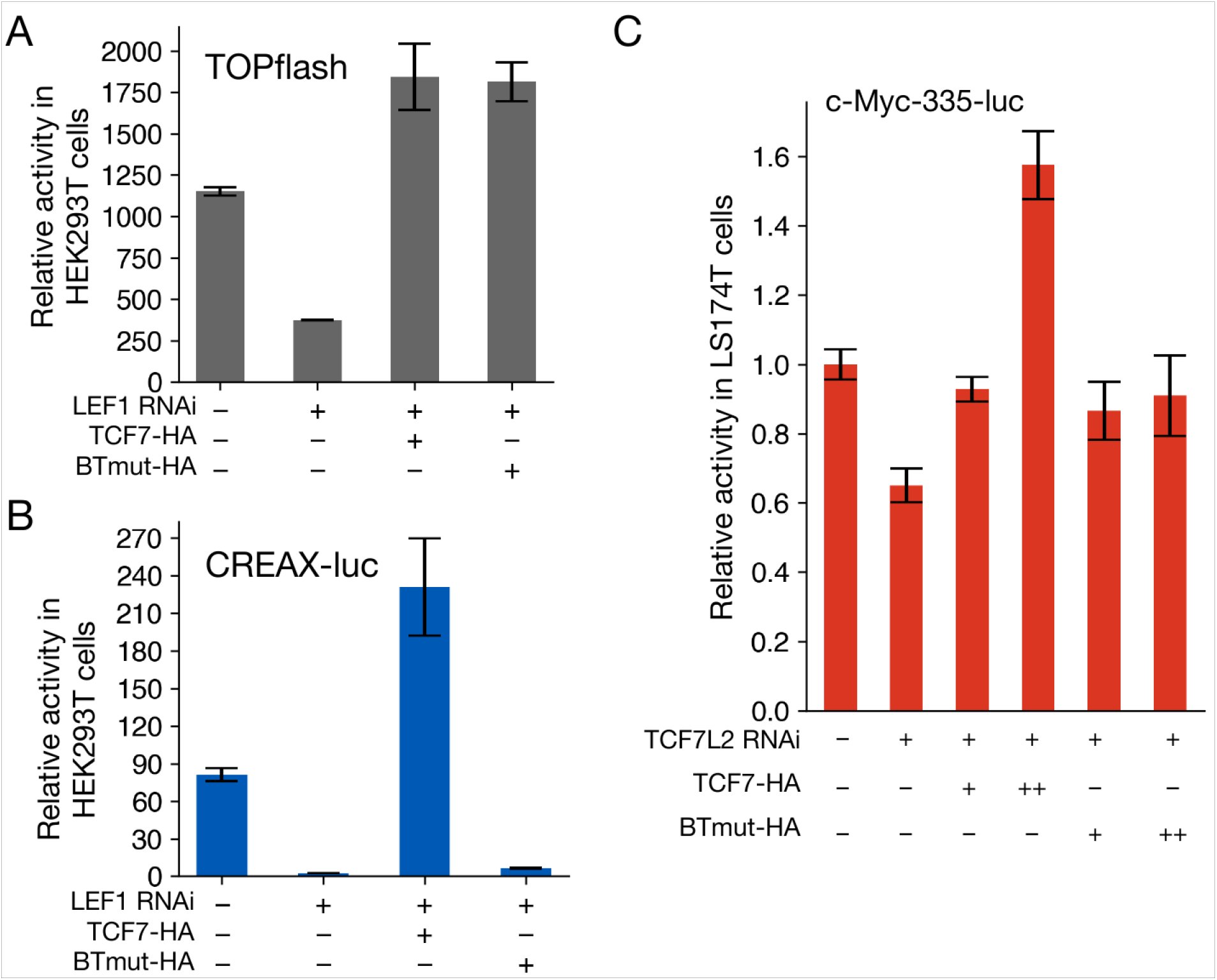
TCF-CDX interactions are important for the activation of natural WREs. A) Luciferase assay showing that TOPflash activity can be driven by both WT TCF7 and BTmut. HEK293T cells were transfected with TOPflash reporter, β-catenin*, and plasmids expressing HA-tagged TCF7, BTmut, and shRNA against LEF1 as indicated. B) Luciferase assay showing that BTmut cannot drive CREAX-luc activity. HEK293T cells were transfected similarly to the TOPflash experiment above. C) Luciferase ability showing a qualitative difference in the ability of TCF7 and BTmut to drive c-Myc-335 reporter expression in LS174T cells. Cells were transfected with the c-Myc-335-luc reporter and the indicated shRNA and protein expressing plasmids. Data are shown as mean ± SD from three replicates (N = 3) for each condition in all panels.

BTmut was also similarly competent for rescuing TOPflash activity (Fig. 4A), strongly suggesting that the BTmut protein still retained its ability to enter the nucleus and bind to DNA and β-catenin. Strikingly, in marked contrast to TOPflash, BTmut overexpression failed to rescue CREAX-luc activity (Fig. 4B). These results support a model of TCF-CDX protein-protein interactions having an essential role in activating CREAX. In LS174T cells, we found that the effect of TCF7L2 RNAi on c-Myc-335 reporter activity could be fully rescued by TCF7 overexpression, while BTmut displayed limited rescuing activity (Fig. 4C). In sum, these experiments provide strong evidence that TCF-CDX interactions play an essential role in activating multiple WREs.

### Identification of TCF/CDX/CAG site clusters in CREAX

One of the key ideas of the TF collective model is that enhancers are bound by a variety of TFs, implying the existence of several distinct, functionally important regulatory motifs in these enhancers (Spitz and Furlong, 2012). We wondered whether there were factors in addition to TCF and CDX proteins that regulated CREAX activity. We sought to identify which portions of the CREAX sequence were important for Wnt-responsive activity by systematic mutagenesis of the CREAX-luc reporter plasmid. First, we looked for TCF-binding sites in the enhancer using the FIMO utility from the MEME suite (Grant et al., 2011) and a list of functionally validated TCF-binding sites (Archbold et al., 2014). Based on this search, we flagged 4 potential TCF-binding sites identified by FIMO with a p-value threshold of < 0.001 and designated these regions T1-T4. We then divided up the remaining regions in the enhancer into blocks of roughly 10 nucleotides each, splitting the 420bp enhancer into 42 blocks. We then generated 42 constructs, each containing mutations in a different block and tested their ability to be activated by β-catenin* in HEK293T cells. In order to compare results across independent experiments, we performed each experiment with the WT CREAX-luc reporter as a control and expressed the fold activation of the mutant constructs as a percentage of the WT reporter.

We found that mutations in the 4 annotated TCF sites reduced the fold activation to at least 55% of the WT reporter. To identify additional regions of regulatory importance, we examined constructs which showed less than 55% of WT activation and those showing more than 100% of WT activity levels (red and blue lines, Fig. 5A). 28 out of 42 constructs fell outside these thresholds. 16 of them showed lower activity upon mutation, suggesting that they contained motifs required for activating the enhancer, while 12 mutant constructs showed higher activity, implying a repressive function (Fig. 5B).

**Fig 5:**
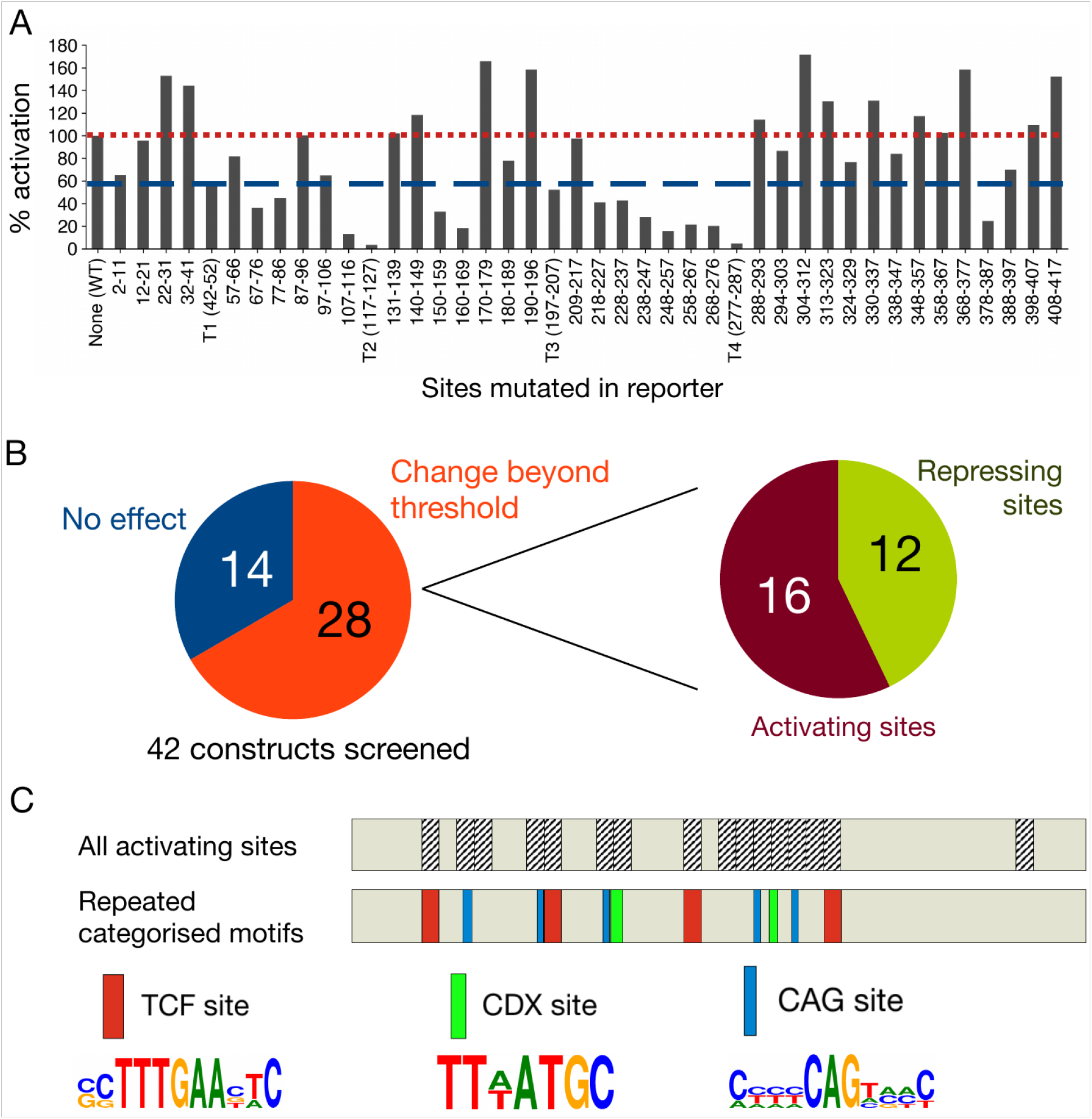
Identification of multiple sets of recurring DNA motifs important for the activation of CREAX by Wnt signalling by a systematic mutagenesis screen. A) Luciferase assay showing fold activation by β-catenin* of CREAX mutant constructs in HEK293T cells. Four TCF-binding sites were identified and mutated (labelled T1-T4), and the remaining nucleotides were mutated as indicated for non-overlapping screening mutagenesis. Mutations which decreased fold activation to at least 55% (blue line) or more than 100% of WT (red line) were considered hits from the screen. Bars represent a mean of two independent experiments (N = 2). B) 28 of the 42 CREAX mutagenesis contructs showed significant changes in activation by β-catenin*. Of the 28, 16 showed reduced activity (suggesting that these regions play a role in enhancer activation) and 12 showed increased activity (suggesting a repressive effect). C) The cartoon depicts the 420 bp CREAX fragment with the activating regions annotated. In addition to TCF binding sites (red), CREAX activity is regulated by multiple CDX binding sites (green) and CAG sites (teal), which are shown with their site logos.

To understand the requirements for the transcriptional activation of CREAX by the Wnt pathway, we examined the 16 sequence blocks containing activating information. In these, we looked for the existence of recurring motifs besides the TCF sites we had already flagged. Among the non-TCF site regions, we found 2 nearly identical sequences (TTTATGC and TTAATGC), which fit the consensus binding site for CDX family proteins (Berger et al., 2008). Additionally, we found five copies of a novel motif centred around the consensus sequence CAG, which we named ‘CAG sites’ (Fig. 5C).

### TCF, CDX, and CAG sites are functionally important for the activity of CREAX and the c-Myc-335 WRE

To confirm the role of TCF, CDX, and CAG sites in CREAX activity, we generated reporter constructs with targeted mutations in the respective sites. Strikingly, we found that abrogating the function of any of the 3 sets of motifs caused a severe reduction in CREAX reporter activation by β-catenin* (Fig. 6A). To confirm that the putative CDX sites we had identified were specifically recognized by CDX protein, we performed gel-shift assays with recombinant GST-CDX1 and a probe containing a TTTATGC sequence. GST-CDX1 caused a gel mobility shift in the biotinylated probe, and the addition of excess unlabelled probe decreased the shift intensity in a manner consistent with competition (Fig. 6B). These results provided support that the sites we identified were indeed CDX sites.

**Fig 6:**
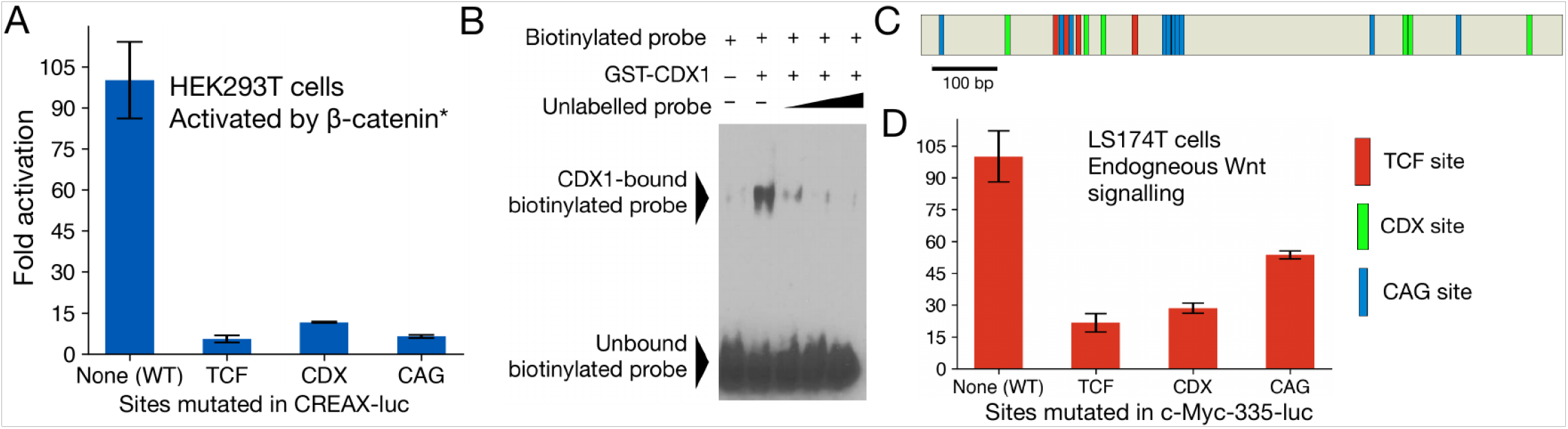
TCF and CDX binding sites and CAG sites are functionally important for CREAX and the c-Myc-335 enhancer. A) Effect of mutating the 4 TCF sites, 2 CDX sites, or the 5 CAG sites on CREAX-luciferase reporter activity in HEK293T cells. Cells were transfected with the indicated reporters and an empty vector or plasmid expressing β-catenin* and the fold activation by β-catenin* was compared. B) CDX1 can bind to the CDX sites in CREAX in vitro. A biotin-labelled probe containing CDX sites from CREAX binds to GST-tagged recombinant CDX1 in an EMSA. An unlabelled probe consisting of these CDX sites competes with the biotinylated probe for binding GST-CDX1. C) Cartoon of the c-Myc-335 enhancer showing TCF binding sites (red), CDX binding sites (green), and CAG sites (teal). D) Luciferase assay showing that the mutation of TCF, CDX, or CAG sites in the c-Myc-335 enhancer decreases its activity in LS174T cells. Cells were transfected with the indicated reporters and their relative activity was analysed by a luciferase assay. Data in (A,D) are shown as mean ± SD from three replicates (N = 3).

Similarly, our examination of the c-Myc-335 enhancer identified 4 potential TCF binding sites, 6 CDX sites, and 10 CAG sites (Fig. 6C). To examine their role in WRE regulation, we constructed reporters with each class of motif mutated. As was the case for CREAX in HEK293T cells, we found that all three mutant reporters displayed a dramatic reduction in activity in LS174T cells (Fig. 6D).

### CAG sites are enriched in regions bound by both TCF7L2 and CDX2

After establishing the functional importance of CAG sites in two WREs, we were interested in whether regulation by TCF, CDX, and CAG sites was a feature of other WREs as well. A previous ChIP-chip study in chromosomes 8, 11, and 12 of LS174T cells had identified 118 loci bound by both TCF7L2 and CDX2 (Verzi et al., 2010). We generated a position-weighted matrix using the 15 CAG sites from CREAX and c-Myc-335 (site logo in Fig. 7A) and used it to look for CAG sites in the 118 doubly-bound loci using the FIMO utility.

**Fig 7:**
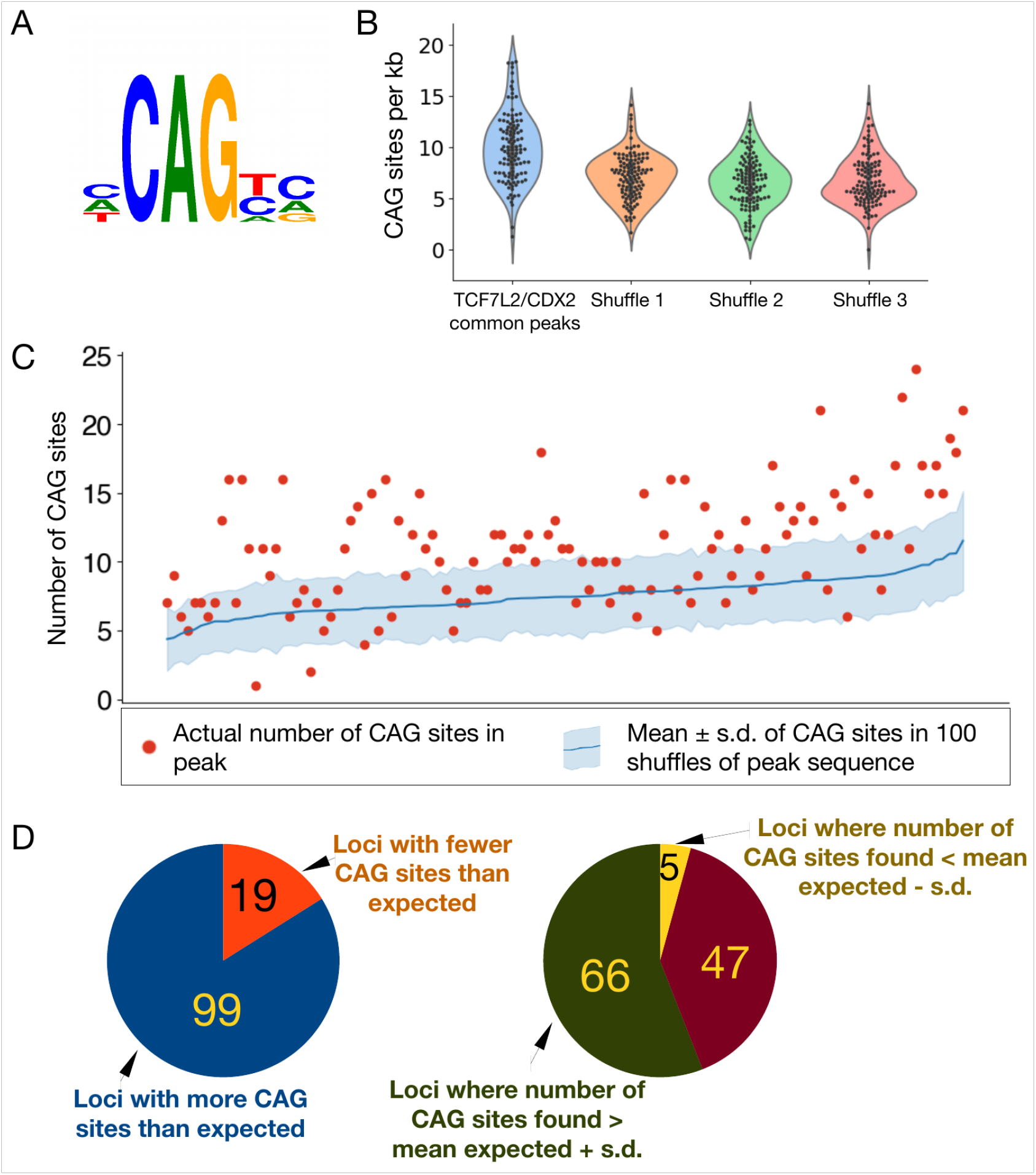
CAG sites are enriched at loci bound by TCF7L2 and CDX2. A) Site logo of functionally validated CAG sites identified in the CREAX and c-Myc-335 WREs. B) Distribution of CAG sites in regions bound by TCF7L2 and CDX2 in colorectal cancer cells from Verzi et al. Each data point shows the number of CAG sites per kb in a single ChIP-chip peak. A search for CAG sites was performed in 3 shuffled versions of each peak’s sequence to see if there was an enrichment of the CAG site motif in these sites. C) CAG sites are enriched in loci bound by TCF7L2 and CDX2. Actual number of CAG sites identified by FIMO are shown in the red scatter plot. The number of CAG sites was calculated in 100 shuffled variants of the sequence of each locus, and the blue line shows the average number of CAG sites in these variants. The shaded region represents the standard deviation in the number of CAG sites. (N = 100) D) 99 (83.9%) of loci bound by both TCF7L2 and CDX2 had more than the expected number of CAG sites. The CAG site count was more than one standard deviation above the mean in 66 of those loci.

The ChIP-chip peaks common to TCF7L2 and CDX2 range in length from 695 to 1594 bp with a mean length of 1090 bp. On average, we found that the common peaks contained 9.93 CAG sites per kb (Fig. 7B). To understand if this represented an enrichment of CAG sites, we compared the number of CAG sites found in each peak to the number that would be expected due to random chance. For each peak, we shuffled the peak’s sequence 100 times to generate comparison sequence of identical length and nucleotide composition. We then calculated the number of CAG sites in each of the 100 shuffled controls and computed the mean and standard deviation to estimate of the number of CAG sites expected from a random distribution (Fig. 7C). Surprisingly, 99/118 (83.9%) common peaks had more CAG sites than expected from chance, with the number of CAG sites in 66 (55.9%) peaks exceeding the expectation value by more than one standard deviation. Only 5 loci contained significantly fewer CAG sites than expected (Fig. 7D). Put together, our analysis suggests that the tripartite TCF/CDX/CAG site regulatory logic could be a general feature of a family of WREs.

### CDX and CAG sites sensitise WREs to Wnt signalling in a cell-type specific manner

Having identified the importance of TCF, CDX, and CAG sites in WREs, we then attempted to generalise our findings using synthetic WREs based on the design principles of CREAX and c-Myc-335. We generated 5 reporters with different combinations of TCF, CDX, and CAG sites. The first reporter contained 3 TCF binding sites alone, similar to the TOPflash reporter corresponding to the conventional model of WREs. The second and third reporters consisted of TCF and either CDX or CAG sites respectively. The fourth reporter contained a combination of CDX and CAG sites without TCF sites, and the fifth contained TCF, CDX, and CAG sites, reflecting the composition of CREAX and c-Myc-335 (Fig. 8A). The spacing between the sites were kept constant to discount any effects caused by changes in relative positions between TF binding sites. We then tested all 5 reporters in 3 cell lines – HEK293T cells, HeLa cells, and LS174T cells. HEK293T and HeLa cells have low basal levels of Wnt signalling, so we compared the fold activation shown by each reporter upon transfection with β-catenin*. Since LS174T cells have a constitutively active Wnt pathway, we compared the relative activity of each reporter to the empty vector (Fig. 8B).

**Fig 8:**
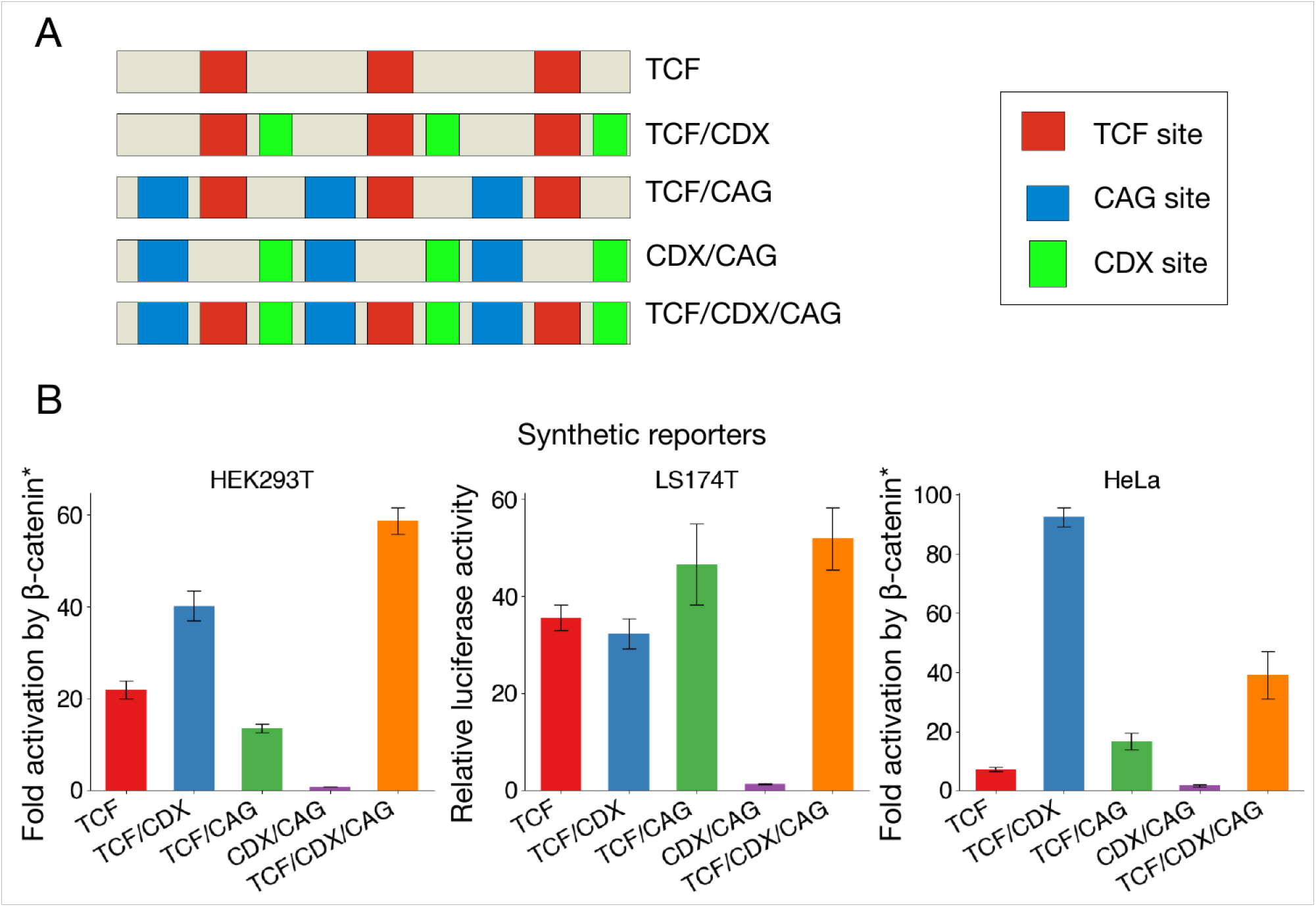
CDX and CAG sites sensitise WREs to Wnt signalling. A) Cartoons of the five synthetic WREs with different combinations of TCF, CDX, and CAG sites. Constructs shown were cloned upstream of a minimal promoter to generate luciferase reporters. B) Relative activity levels of the synthetic WREs in HEK293T, LS174T, and HeLa cells. HEK293T and HeLa cells were transfected with indicated reporters and an empty vector or β-catenin* expression plasmid. Fold activation by β-catenin* was compared. LS174T cells were transfected with indicated reporters, and activity was normalised to that of the empty vector with only the minimal promoter. Each bar represents mean ± SD from three replicates (N = 3)

These experiments yielded three main conclusions. Firstly, we found that the TCF/CDX/CAG construct more active than the construct containing only TCF sites in all 3 cell lines. From this we concluded that CDX and CAG sites play a general role in sensitising WREs to Wnt signalling. Secondly, the construct lacking TCF sites and containing only CDX and CAG sites did not show Wnt-responsive activity, supporting a model of them supporting WRE activation through TCFs. Thirdly, the best-performing WREs were different in each cell type – while the TCF/CDX/CAG was the clear winner in HEK293T cells, it showed a similar level of activity as the TCF/CAG reporter in LS174T cells. Similarly, while both the TCF/CAG and TCF/CDX/CAG were more sensitive than the TCF-only reporter in HeLa cells, the TCF/CDX construct was more than twice as sensitive as the TCF/CDX/CAG construct. When looked at simultaneously, the contributions of TCF, CDX, and CAG sites to WRE activity across cell types paint a picture of a TF collective containing TCF and CDX proteins whose activity is modulated by additional cell type specific factors.

## Discussion

The question of what makes certain regions of the genome act as enhancers has long been of interest in the field of gene regulation. It is widely accepted that enhancers contain clusters of binding sites for multiple TFs. However, understanding the “binding site grammar” of enhancers – the rules governing the composition, distribution, and orientation of TF binding sites that allow them to drive expression under specific conditions – remains a challenge (Lambert et al., 2018; Wasserman and Sandelin, 2004). The problem of enhancer composition is typically discussed using two overarching models, the enhanceosome and the flexible billboard. The enhanceosome is best exemplified by the *Interferon-β* promoter (Panne et al., 2007; Thanos and Maniatis, 1995) and models enhancers as consisting of specific TF binding sites organised in a specific order and orientation. The flexible billboard model portrays enhancers as collections of TF binding sites which vary in their composition and arrangement. Enhancer activation occurs when a sufficient number of TF binding sites are occupied (Arnosti and Kulkarni, 2005). While there is experimental evidence of some constraint in binding site order and orientation on some enhancers (Farley et al., 2016; Jolma et al., 2015; Swanson et al., 2010), other studies support the flexible billboard view (Junion et al., 2012; Smith et al., 2013). The TF collective model adds a further level of complexity to the billboard model by invoking protein-protein interactions between TFs as an additional means of recruiting TFs to enhancers (Spitz and Furlong, 2012). While computational efforts to understand this enhancer grammar have had some success incorporating aspects of all three aforementioned models (Chen and Capra, 2020), our results with WREs demonstrate the need for additional bottom up experimental approaches to understand the logic of enhancer composition.

Here, we report on two WREs whose binding site grammar is consistent with the billboard rubric of enhancer function. A combination of ChIP and binding site mutagenesis indicate that the CREAX and c-Myc-335 WREs are direct targets of TCF and CDX TFs (Figs. 2,5). Systematic mutagenesis of CREAX revealed the presence of multiple copies of another motif, which we have termed CAG sites (Fig. 5). These CAG sites are required for Wnt responsiveness of the CREAX and c-Myc-335 WREs (Fig. 6). CAG motifs are also enriched in the vast majority of regions bound by TCF7L2 and CDX2 in a colorectal cell line (Fig. 7). While TCF/CDX/CAG site-based regulation appears to define a significant class of WREs, the arrangement of the binding sites is highly variable. For example, in CREAX, the TCF and CAG sites are roughly evenly spread out, while in the c-Myc-335 enhancer, 3 out of 4 TCF sites and 5 of the CAG sites form separate clusters (Figs. 5C, 6C). In the 118 sites bound by TCFL2 and CDX2, there is also a great flexibility of motif composition, with the density of CAG sites ranging from 1.3 – 21.3 sites per kb. While all these sites have not been functionally tested, the systematic analysis of CREAX supports a model where all the TCF, CDX and CAG sites contribute to activation by Wnt signalling (Fig. 5A).

In addition to supporting the billboard model for WRE function, our data demonstrating the importance of TCF-CDX physical interactions also fits with the TF collective vision of enhancer activation. Our work extends previous findings demonstrating an interaction between the basic tail of TCFs and the homeodomain of CDXs (Béland et al., 2004a). We found that mutation of two residues in the basic tail of TCF7 (BTmut), resulting in a decreased binding to CDX1, did not affect its ability to activate a TCF-site reporter. However, BTmut was dramatically crippled in supporting CREAX activation and showed a significant defect in regulating c-Myc-335 (Fig. 4). Our data provide compelling support for the importance of TCF-CDX interactions in WRE activation, but the precise mechanism remains to be elucidated. A previous study identified the recruitment of CDX1 and LEF1 to CDX1 regulatory DNA, and the absence of CDX binding sites in this sequence led the authors to surmise that TCFs could recruit CDX proteins to DNA (Béland et al., 2004a). Alternatively, CDXs have been reported to act as pioneer factors, making chromatin more accessible to binding by other TFs (Kumar et al., 2019; Verzi et al., 2013). This raises the possibility that the essential CDX-TCF interaction we have identified occurs after CDX-induced chromatin opening. Alternatively, complex formation with pioneer factors could allow certain WREs to be activated sooner than others and provide a means for temporally regulating the Wnt response.

Our findings on how CDXs can directly co-regulate WREs could provide mechanistic insight into previous findings in vertebrate development. Wnt/β-catenin signalling, TCFs and CDXs have all been demonstrated to be important for axial patterning in mammalian development, and specifying posterior/caudal cell fates (Galceran et al., 1999; Lohnes, 2003; Shimizu et al., 2005). CDXs are transcriptional targets of Wnt signalling (Pilon et al., 2006, 2007) suggesting a hierarchy of action. However, consistent with our results, there is also evidence that CDXs and the Wnt pathway function at the same level of the axial patterning hierarchy (Amin et al., 2016; Young et al., 2009). This suggests that TCF/β-catenin and CDX may also directly act on enhancers controlling posterior fate genes. Wnt signalling and CDXs are also known to intersect in other contexts such as the haematopoietic system (Lengerke et al., 2008) and primordial germ cells (Bialecka et al., 2012). Mice engineered to express the BTmut variant could be a powerful tool to test the importance of WRE activation by CDX proteins in these and other contexts.

Our synthetic enhancer experiments demonstrated that CDX and CAG sites contribute to WRE function across cell types (Fig. 7). The addition of CDX sites increased WRE activity in HEK293T and HeLa cells, while it had no effect in LS174T cells. One explanation for this observation could be that while TCF and CDX TFs directly bind to these WREs in all 3 cell types, the TF collectives regulating these WREs contain additional cell type-specific TFs which contribute to enhancer activity without binding to DNA. A similar model has been proposed for TBX3 regulation of Wnt targets in mammalian limb development (Zimmerli et al., 2020). Similarly, the addition of CAG sites increases WRE activity in LS174T and HeLa cells but decreases it in HEK293T cells. Since the identity of the CAG site binding protein is still unknown, this could imply the existence of multiple TFs capable of binding CAG sites expressed at different levels in various cell types. In addition to the cis-regulatory information that recruits TFs to an enhancer, the trans-regulatory environment of the cell can dictate the identity of the TF collective which binds to the enhancer. Transcriptional regulation supplementing the direct binding of TFs to DNA provides additional control over enhancer activity to produce the exquisite spatio-temporal specificity of transcription seen throughout animal development.

The systematic mutagenesis of the CREAX enhancer provides a dramatic illustration that naturally occurring WREs consist of more than clusters of TCF binding sites. Our identification of CDX and CAG sites in WREs refines our understanding of the binding site grammar underlying some WREs. It is important to note that the identified TCF, CDX and CAG sites only account for 11 of the 16 regions required for activation of CREAX (Fig. 5C). In addition, there are 12 regions with repressive activity. These functional regions don’t contain consensus AP-1 sites, which have been implicated in WRE function (Yochum et al., 2008). Nor do they contain the repressive motif recently identified in several WREs (Kim et al., 2017). Future experiments will explore how these sequences interact with the TCF/CDX/CAG cassette, to more fully understand the multiple layers of regulation in WREs.

## Materials and Methods

### Cell culture and transfection

All cell lines were grown at 37°C and 5% CO_2_. HEK293T (ATCC, CRL-3216) and HeLa (ATCC, CCL-2) were grown in DMEM (Dulbecco’s Modified Eagle Medium, Gibco, 11995065) supplemented with 10% foetal bovine serum (FBS) and penicillin/streptomycin/glutamine (PSG, Gibco, 10378016). LS174T cells (ATCC, CL-188) were cultured in MEM (Minimum Essential Medium, Gibco, 11095080) containing 10% FBS and PSG. Cells were transfected using Lipofectamine 2000 (Invitrogen, 11668030) or PEI MAX (Polysciences, 24765-1) according to manufacturer instructions.

### Plasmids

The CREAX fragment was amplified from human genomic DNA with the following primers: 5’-CTTGCTCGAGGGAACCCGCTGAATGGCTGG-3’, 5’-ACTCAGATCTCAACACAGCGCTCCTGTCCA-3’, and cloned into upstream of the minimal promoter in pGL4.23 (Promega, E841A) with XhoI and BglII. Mutations were made in the CREAX reporter using the Quickchange II Kit (Stratagene, 200518). Details of the mutations are shown in supplementary table 1. The TOPflash and Ax2 reporters (Leung et al., 2002) along with the β-catenin* (S33Y mutation, cloned into pcDNA3.1) expression plasmid were a kind gift from Dr. Eric Fearon, University of Michigan. Expression constructs for shRNAs were generated by cloning in the appropriate oligonucleotides into the pSUPER vector (OligoEngine, VEC-PBS-0002) according to the manufacturer’s instructions. Targeting sequences for the shRNA constructs are listed in supplementary table 2.

The TCF7 open reading frame (ORF) was amplified with PCR (primers: 5’-GAATTCGAGCACTGTCATCGGAAGGAAC-3’ and 5’-GGTACCATGCCGCAGCTGGACTCC-3) and cloned into pcDNA3.1 with KpnI and EcoRI. The CDX1 ORF was similarly amplified and cloned (primers: 5’-GGTACCATGTATGTGGGCTATGTGCTGGAC-3’ and 5’-GAATTCTGGCAGAAACTCCTCTTTCACAG-3’). The HA tag was generated by annealing two oligonucleotides (5’-AATTCTACCCATACGATGTTCCAGATTACGCTTACCCATACGATGTTCCAGATTACGCTTACCCATACGATGTTCCAGATTACGCTTAAT-3’ and 5’-CTAGATTAAGCGTAATCTGGAACATCGTATGGGTAAGCGTAATCTGGAACATCGTATGGGTAAGCGTAATCTGGAACATCGTATGGGTAG-3’) which were then cloned into protein expression plasmids using EcoRI and XbaI. Flag tags were added in a similar manner using different oligos (5’-AATTCGACTACAAGGATGACGATGACAAAGACTACAAGGATGACGATGACAAAGACTACAAGGATGACGATGACAAATAAT-3’ and 5’-CTAGATTATTTGTCATCGTCATCCTTGTAGTCTTTGTCATCGTCATCCTTGTAGTCTTTGTCATCGTCATCCTTGTAGTCG-3’). GST-CDX1 was generated by PCR cloning the CDX1 ORF into pGEX-6P-1 (Cytiva, 28-9546-48) using the following primers: 5’-CGCGGATCCATGTGCTGGACAAGGATTCG-3’ and 5’-ATAGTTTAGCGGCCGCTTATGGCAGAAACTCCTCTTTCACAG-3. His-TCF7 was PCR cloned into pET52b (Millipore, 71554-3) using the primers 5’-CGGGGTACCTCATGCCGCAGCTGGACTCC-3’ and 5’-ATAGTTTAGCGGCCGCGAGCACTGTCATCGGAAGGAAC-3’. Site-directed mutagenesis was performed using the primers 5’-GAAGAGGCGGTCGGAGGAAGAGCACCAAGAATCCAC-3’ and 5’-GTGGATTCTTGGTGCTCTTCCTCCGACCGCCTCTTC-3’ to generate HA- and His-tagged basic tail mutant versions of TCF7.

The synthetic TCF/CDX/CAG reporter was created by annealing the following oligos: 5’-GATCTCCAACAGTCACGGTACCTTTGATCTTGTAGTTTATGCGTACCAACAGTCACGGTACCTTTGATCTTGTAGTTTATGCGTACCAACAGTCACGGTACCTTTGATCTTGTAGTTTATGCC-3’ and 5’-TCGAGGCATAAACTACAAGATCAAAGGTACCGTGACTGTTGGTACGCATAAACTACAAGATCAAAGGTACCGTGACTGTTGGTACGCATAAACTACAAGATCAAAGGTACCGTGACTGTTGGA-3’, and cloning into pGL4.23 with BglII and XhoI. The other synthetic reporters were generated by site-directed mutagenesis of this reporter using primers shown in supplementary table 3. Directed mutations made to the CREAX and c-Myc-335 reporters are listed in supplementary table 4.

### Luciferase assays and analysis

Luciferase assays were performed with firefly-luciferase reporter plasmids and either β-galactosidase (LacZ) or renilla luciferase as an internal transfection control. Activities of LacZ and luciferase were measured using the Tropix Galacto-Star (Applied Biosystems, T1056) and Tropix Luc-Screen systems (Applied Bisosystems, T1035). Experiments with firefly and renilla luciferase were assayed using Promega’s Dual-luciferase reporter assay system (Promega, E1910) as per the manufacturer’s protocol. For each well assayed, the firefly/LacZ or firefly/renilla ratio was used as the measure of luciferase activity. All luciferase assays were done in triplicates, except for the mutagenesis screen in Fig. 2A, which was done with duplicates of each data point. The mean and standard deviation of the firefly/LacZ or firefly/renilla ratios were calculated. Data from the luciferase assays are expressed in terms of relative luciferase activity (RLA) or fold activation. To calculate RLA, a basal condition was selected (specified in each figure). The mean and standard deviation of the ratios of all conditions were then proportionally expressed in terms of the mean of the ratios of the basal conditions. Fold activation was calculated as the quotient of the luciferase activity under the specified Wnt-activated condition to the activity without Wnt signalling. Both Wnt on and Wnt o ff conditions were done in triplicate with appropriate error propagation used to calculate the standard deviation in the quotient.

### Measurement of transcript levels by RT-qPCR

Cells were lysed with TRIzol (Invitrogen, 15596026) and RNA extraction was performed using the Rneasy mini kit (Qiagen, 74104). Reverse transcription was performed using SuperScript III Reverse Transcriptase (Invitrogen, 18080093). Quantitative PCR was then performed on a CFX Connect Real-Time PCR Detection System (Bio-Rad, 1855201) using the Power SYBR Green PCR Master Mix (Applied Biosystems, 4368577). The mean and standard deviation of transcript levels were quantified using the Pffafl method (Pfaffl, 2001) with *Axin2* transcript levels normalised to *G6PD* transcript levels.

### Gel shift assay

EMSA was performed as described previously (Chang et al., 2008) using a 6% native gel. GST-tagged CDX1 and 20 fmol biotinylated DNA probe (5’-Bio-GGCCAACAGTCACGGTACCTTTGATCTTGTAGTTTATGCGTACCAACAGTCACGGTACCTTTGATCTTG TAGTTTATGCGT-3’) were incubated with 50μg/ml poly (dI-dC), 0.05% NP-40, 5 mM MgCl2 and 2μl of 50% glycerol in the presence of binding buffer (10 mM Tris–HCl, pH 7.5, 50 mM KCl, 1 mM DTT) for 5 min on ice and 20 min at room temperature. For competition assays, unlabelled CDX binding probe (5’-TCTTGTAGTTTATGCGTACGTAGTTTATGCGTACC-3’) was incubated with the reaction mixture containing protein for 10 min prior to adding the labelled probe.

### CAG site enrichment analysis

A list of CAG sites identified in the CREAX and *c-Myc* WREs was generated and converted into a MEME motif file using the sites2meme utility from the MEME suite (Bailey et al., 2009). To calculate an appropriate p-value threshold to search for CAG sites across the genome, the FIMO utility (Grant et al., 2011) was run with the CAG site motif file to identify CAG sites in the c-Myc-335 WRE. The CAG site with the highest p-value in the c-Myc-335 WRE had a p-value of 0.003611, and this was set as the threshold.

Data files analysed came from a previously published study examining the binding of TCF7L2 and CDX2 using ChIP-chip (Verzi et al., 2010). Raw data from the original study are publicly available online (GEO accession no. GSE22572). Files containing the coordinates of regions bound by TCF7L2 and CDX2 were kindly provided to us by Dr. Michael P. Verzi. The bedtools intersect program (Quinlan and Hall, 2010) was used with the ‘-wa’ option to generate a BED file containing the coordinates of the overlapping TCF7L2 and CDX2 peaks. The twoBitToFa utility from the UCSC genome browser (Kent et al., 2002) was then used to obtain the sequences of each of the regions bound by TCF7L2 and CDX2. FIMO was then run to identify CAG sites using the following command: “fimo --thresh 0.003611 CAG-sites-meme.txt tcf7l2-cdx2-common.fasta”. Shuffled versions of these sequences were generated using the fasta-shuffle-letters utility from the MEME suite: “fasta-shuffle-letters -dna tcf7l2-cdx2-common.fasta LS174T-shuffle-001.fasta”. This command was run 100 times to generate 100 different shuffles. The number of CAG sites in each of the original and shuffled peaks were tabulated and the number of sites per kb was calculated.

### Antibodies

Western blots were probed with the following antibodies: Anti-Flag-HRP (Sigma-Aldrich, A8592), Anti-HA (Roche, 11867423001), Anti-His (Cytiva, 27-4710-01), and Anti-GST (Invitrogen, A-5800). Anti-Flag immunoprecipitations were carried out using magnetic beads pre-conjugated with the antibody (Millipore, M8823). Anti-TCF7L2 (Cell Signaling Technology, 2569S) antibodies were also used for ChIP.

### Generation of a Flag-tagged cell line using CRISPR/Cas9

A guide RNA targeting a site immediately downstream of the human CDX1 stop codon (atgcccaccctgtgccccgg) was designed using CRISPOR (Concordet and Haeussler, 2018). This was then cloned into PX458, a plasmid expressing Cas9 protein (Ran et al., 2013). pSpCas9(BB)-2A-GFP (PX458) was a gift from Feng Zhang (Addgene plasmid # 48138). The homology-directed repair template was designed based on the pFETCh_Donor plasmid from the CETCh-seq protocol (Savic et al., 2015). pFETCh_Donor (EMM0021) was a gift from Eric Mendenhall & Richard M. Myers (Addgene plasmid # 63934). Since HEK293T cells are G-418 resistant, the neomycin resistance gene in the plasmid was changed to puromycin resistance. The puromycin resistance gene from sgOpti (Fulco et al., 2016) was PCR amplified using the following primers – 5’-atcgtttcgcatgacagaatacaaaccaaccgtgc-3’ and 5’-gtataagaagacatgatgttcgaatcaagcaccaggttttcgtgtc-3’. sgOpti was a gift from Eric Lander & David Sabatini (Addgene plasmid # 85681). pFETCh_Donor was digested with AleI-v2 and BstBI (NEB). The 3x-Flag-P2A cassette was amplified from pFETCh_Donor (primers: 5’-atactctgtaatcctactcaataaacgtgtcacg-3’, 5’-attctgtcatgcgaaacgatccaggtccagggttc-3’). The fragments were then joined by Gibson assembly to create pFETCh_Puro, in which the neomycin resistance cassette was replaced with puromycin resistance. HEK293T genomic DNA was purified using the Dneasy Blood & Tissue kit (Qiagen, 69504). Homology arms were amplified using the following primers – arm 1 with 5’-ctgacgtcgacggatcgggaTCCCCGACCTGCAGCCCA-3’ and 5’-CCGGAACCTCCTCCGCTC-3’ and arm 2 with 5’-ttcgaacatcCTCGGGTGCTGGGAGTGT-3’ and 5’-actgtgctggatatctgcagGCTCTGCTTGGTCCGAATAAAG-3’. The 3xFlag-P2A-Puromycin construct was amplified using the following primers: 5’-gggagcggaggaggttccggTGGAGGTGGTTCTGGAGATTAC-3’ and 5’-agcacccgagGATGTTCGAATCAAGCACC-3’. The pcDNA3.1 plasmid (Invitrogen, V79020) was digested with EcoRI-HF and BglII (NEB) and the homology arms and flag-tagging constructs were assembled into the plasmid using Gibson assembly to generate the repair template for genome editing.

HEK293T cells were co-transfected with the Cas9/gRNA expressing plasmid and the repair template. Puromycin selection was performed 48 hours post-transfection using 500 ng/mL puromycin. Over 150 distinct puromycin-resistant colonies of cells could be seen, and these were allowed to grow under selection to create a polyclonal cell population. Cells were subsequently grown with 250 ng/mL puromycin to maintain selection. Genomic DNA was purified using the Dneasy blood & tissue kit to confirm the presence of the edits. PCRs were set up with primers specific to the Flag construct (5’-caatgcctgtgaaagaggagtttctg-3’ and 5’-CAATCTTTCATAAAAAGGCAGATTTCG-3’) and primers flanking the insertion site, which would generate a 450 bp amplicon from the unedited locus (5’-tcttgcactctctctttcactctctcc-3’ and 5’-tttatccaacaggcttactgcacagat-3’). The sequence of the edited locus was confirmed by Sanger sequencing the product from the Flag-specific PCR.

### Chromatin immunoprecipitation (ChIP)

Cells were fixed in 0.75% formaldehyde at room temperature for 10 minutes. They were then incubated for 5 minutes with 125 mM glycine to quench the formaldehyde. Following this, they were rinsed thrice with PBS and resuspended in RIPA buffer (Sigma-Aldrich, R0278). The cell suspension was sonicated using a Covaris M220 focused ultrasonicator. Cell debris was pelleted by high-speed centrifugation. One fraction of the supernatant was saved as the input fraction and the rest was used for the immunoprecipitation (IP). The cell lysate was pre-cleared with 30 μ L Protein A agarose beads (Millipore, 16-156) for 1 hour at 4°C and then incubated overnight on a rotor with antibody coated beads. The beads were then washed once in low salt buffer (0.1% SDS, 1% Triton X-100, 2 mM EDTA, 20 mM Tris-HCl pH 8.0, 150 mM NaCl), high salt buffer (0.1% SDS, 1% Triton X-100, 2 mM EDTA, 20 mM Tris-HCl pH 8.0, 500 mM NaCl), and LiCl wash buffer (0.25 M LiCl, 1% NP-40, 1% sodium deoxycholate, 1 mM EDTA, 10 mM Tris-HCl pH 8.0). DNA was eluted by gentle shaking in elution buffer (1% SDS, 100 mM NaHCO 3) for 15 minutes. Decrosslinking was performed by adding NaCl to a final concentration of 0.2 M. RNA was digested with Rnase A (Invitrogen, 12091021) followed by treatment with Proteinase K (Sigma-Aldrich, 3115879001). DNA fragments from the input and IP fractions were purified using a PCR purification kit (Qiagen, 28104) and measured with qPCR.

Three sets of primers were used for qPCR – one targeting the CREAX locus (5’-TCCATAGCCAACAGTCACGC-3’ and 5’-GCAATCCTGCCAGCAATCTC-3’) and two targeting flanking regions located 1-2 kb away from CREAX in the 5’ and 3’ directions (Flank 1: 5’-CGTCATCCTGCAACAAGCTG-3’ and 5’-TCTCCATCCACCCTGACCTT-3’; Flank 2: 5’-CTGGGTTGGGGACCACATTT-3’ and 5’-TGTCATTGCCAGCATCACCA-3’). Quantitative PCR was then performed on a CFX Connect Real-Time PCR Detection System (Bio-Rad, 1855201) using the Power SYBR Green PCR Master Mix (Applied Biosystems, 4368577). The percent input was determined for each of 3 replicates and the mean and standard deviation were calculated.

### Nuclear extract preparation and co-immunoprecipitation

Cells were rinsed with ice-cold PBS and then collected into 1 mL of cold PBS. Cells were pelleted and resuspended with 400 μL buffer A (10mM HEPES pH7.9, 1.5mM MgCl2, 10mM KCl, 0.5mM DTT, 0.5% NP-40) supplemented with a protease inhibitor cocktail (Roche, 11873580001). This suspension was incubated on ice for 15 minutes followed by vigorous vortexing for 10 seconds to lyse cell membranes. Nuclei were pelleted by centrifugation. The pellet was resuspended in 100μl buffer B (20mM HEPES pH7.9, 25% Glycerol, 420mM NaCl, 1.5mM MgCl2, 0.2mM EDTA, 0.5mM DTT) supplemented with protease inhibitor cocktail on ice for 20 minutes to lyse nuclei. Cell debris was removed by high-speed centrifugation and the supernatant was used for co-immunoprecipitation.

Three-fold diluted nuclear extracts were pre-cleared with 50 μL Protein-G agarose beads (Millipore, 16-266) for 1 hour at 4°C and then incubated overnight on a rotor with 3 μL anti-FLAG antibody (Sigma, F3165). Each reaction was incubated with Protein-G agarose beads (50 μL) for one more hour. Beads were washed four times with 3-fold diluted buffer B. Proteins were eluted in 2x SDS sample buffer (0.125 M Tris-HCl pH 6.8, 4% (w/v) SDS, 20% (v/v) Glycerol, and 0.01% (w/v) Bromophenol blue) and analysed using western blotting.

### Western blotting

Cell lysates or protein samples were lysed and denatured in hot 2x SDS sample buffer. Protein samples were separated by SDS-PAGE and transferred onto a PVDF membrane (Bio-Rad, 162-0177) and blocked in 5% bovine serum albumin (BSA, Dot Scientific, DSA30075-100). Protein blots were incubated with the appropriate concentration of primary antibody diluted in 5% BSA overnight at 4°C. They were then washed thrice with Tris-buffered saline containing 1% Tween-20 (TBS-T). Blots were incubated with the secondary antibody diluted in 5% BSA for 1 hour. After washing thrice in TBS-T, they were developed using a chemiluminescence substrate (Pierce, 32109) and imaged in a LI-COR Odyssey CLx imager. Images were processed using the GNU Image Manipulation Program.

### Recombinant protein expression and purification

Plasmids expressing GST- and His-tagged proteins were transformed into BL21(DE3) competent cells (Thermo Scientific, EC0114) and grown in LB media at 37°C. Protein expression was induced by the addition of IPTG when the OD reached 0.6. After 3-4 hours, cells were collected by centrifugation and resuspended in the appropriate lysis buffer. Cells were lysed using sonication and proteins were purified using standard protocols (Harper and Speicher, 2011; Nallamsetty and Waugh, 2007).

### GST pull-down assays

5 μg each of the recombinant GST-tagged bait and His-tagged prey proteins were incubated for 1.5 hours in 200 μL of pull-down buffer (20mM Tris-HCl pH7.6, 150mM NaCl, 1%Triton X-100) at 4°C. Glutathione Fast Flow Sepharose 4 beads (GE Healthcare, 17-5132-01) that had been pre-washed in binding buffer were then added to the binding reaction and it was incubated in a rotor for 2 hours at 4°C. The supernatant was then removed and the beads were washed 5 times with 600 μL of binding buffer. Proteins were then eluted using 20 μL of 2x SDS sample buffer and analysed using western blotting.

## Acknowledgements

This work was supported by grants from the US National Institute of Health (R01 GM108468) to K.M.C. Support was also provided by a University of Michigan Rogel Cancer Center Research Grant and a University of Michigan M-cubed award to K.M.C. A.B.R was supported by the Biosciences fellowship from the University of Michigan Department of Molecular, Cellular, and Developmental Biology, a One-Term Dissertation Fellowship and a Graduate Student Research Grant from the Rackham Graduate School, University of Michigan

## Author contributions

LC performed the initial CREAX experiments, the mutagenesis screen, designed synthetic reporters, and characterised the BTmut variant. PB characterised c-Myc-335 and generated the site mutants. ABR performed ChIP experiments, CAG site analyses, c-Myc-335 rescue experiments, and wrote the manuscript. KMC supervised the research and critically revised the manuscript.

## Conflict of interest

The authors declare that they have no conflicts of interest.

---

**Supplementary table 1.**
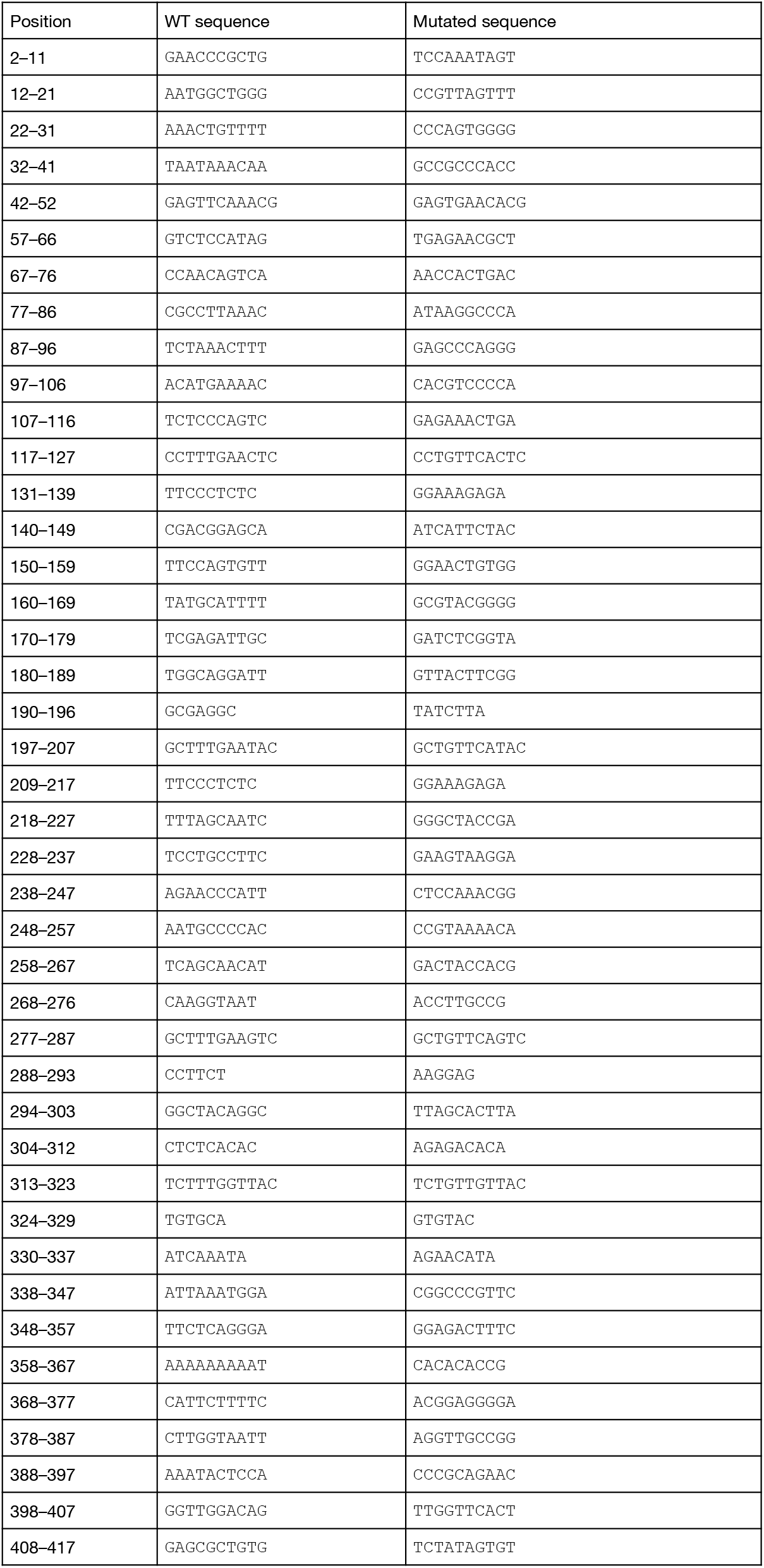
Mutations made in CREAX screening mutagenesis constructs.

**Supplementary table 2.**
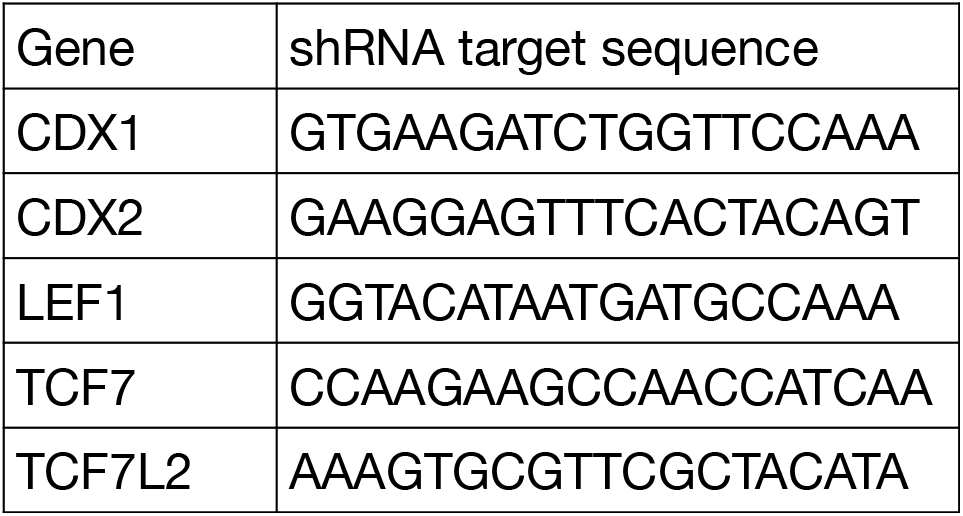
Targeting sequences of the shRNA constructs used in this study.

**Supplementary table 3.**
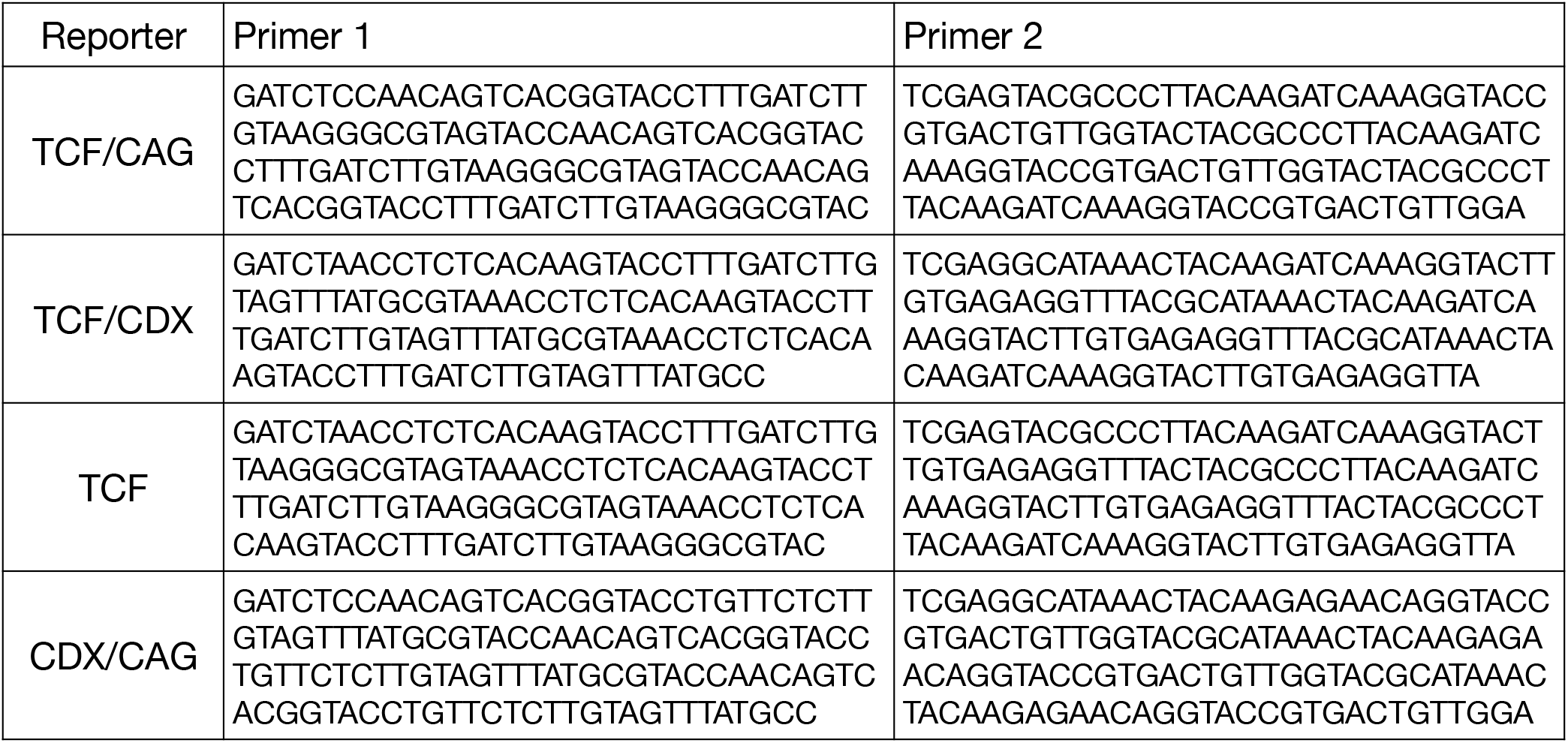
Primers used to generate synthetic reporters.

**Supplementary table 4.**
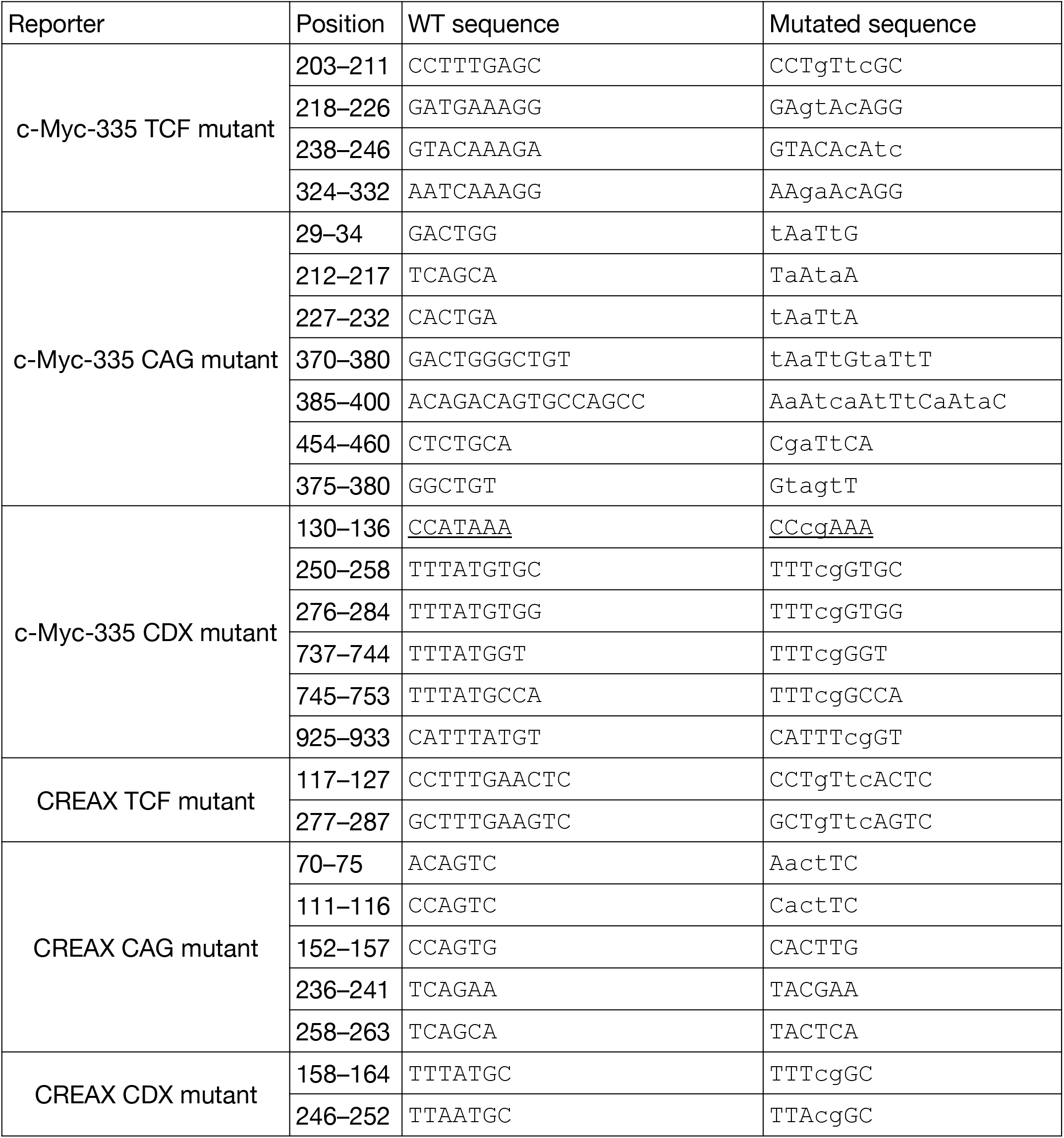
Directed mutations in TCF, CDX, and CAG sites of CREAX and c-Myc-335 reporters.

## Notes

### Competing Interest Statement

The authors have declared no competing interest.

### Summary of Updates

Added author contributions, modified title, added sample size and statistical information to figure legends

## References

Amin, S., Neijts, R., Simmini, S., van Rooijen, C., Tan, S.C., Kester, L., van Oudenaarden, A., Creyghton, M.P., and Deschamps, J. (2016). Cdx and T Brachyury Co-activate Growth Signaling in the Embryonic Axial Progenitor Niche. Cell Reports 17, 3165–3177.

Archbold, H.C., Yang, Y.X., Chen, L., and Cadigan, K.M. (2012). How do they do Wnt they do?: regulation of transcription by the Wnt/β-catenin pathway. Acta Physiologica 204, 74–109.

Archbold, H.C., Broussard, C., Chang, M.V., and Cadigan, K.M. (2014). Bipartite Recognition of DNA by TCF/Pangolin Is Remarkably Flexible and Contributes to Transcriptional Responsiveness and Tissue Specificity of Wingless Signaling. PLOS Genetics 10, e1004591.

Arnosti, D.N., and Kulkarni, M.M. (2005). Transcriptional enhancers: Intelligent enhanceosomes or flexible billboards? Journal of Cellular Biochemistry 94, 890–898.

Atcha, F.A., Syed, A., Wu, B., Hoverter, N.P., Yokoyama, N.N., Ting, J.-H.T., Munguia, J.E., Mangalam, H.J., Marsh, J.L., and Waterman, M.L. (2007). A Unique DNA Binding Domain Converts T-Cell Factors into Strong Wnt Effectors. Molecular and Cellular Biology 27, 8352–8363.

Bailey, T.L., Boden, M., Buske, F.A., Frith, M., Grant, C.E., Clementi, L., Ren, J., Li, W.W., and Noble, W.S. (2009). MEME Suite: tools for motif discovery and searching. Nucleic Acids Res 37, W202–W208.

Barolo, S. (2006). Transgenic Wnt/TCF pathway reporters: all you need is Lef? Oncogene 25, 7505–7511.

Béland, M., Pilon, N., Houle, M., Oh, K., Sylvestre, J.-R., Prinos, P., and Lohnes, D. (2004a). Cdx1 Autoregulation Is Governed by a Novel Cdx1-LEF1 Transcription Complex. Molecular and Cellular Biology 24, 5028–5038.

Béland, M., Pilon, N., Houle, M., Oh, K., Sylvestre, J.-R., Prinos, P., and Lohnes, D. (2004b). Cdx1 Autoregulation Is Governed by a Novel Cdx1-LEF1 Transcription Complex. Mol. Cell. Biol. 24, 5028–5038.

Berger, M.F., Badis, G., Gehrke, A.R., Talukder, S., Philippakis, A.A., Peña-Castillo, L., Alleyne, T.M., Mnaimneh, S., Botvinnik, O.B., Chan, E.T., et al. (2008). Variation in Homeodomain DNA Binding Revealed by High-Resolution Analysis of Sequence Preferences. Cell 133, 1266–1276.

Bhambhani, C., and Cadigan, K.M. (2014). Finding a Needle in a Genomic Haystack. In Wnt Signaling in Development and Disease, S. Hoppler, and R.T. Moon, eds. (John Wiley & Sons, Inc), pp. 73–87.

Bhambhani, C., Ravindranath, A.J., Mentink, R.A., Chang, M.V., Betist, M.C., Yang, Y.X., Koushika, S.P., Korswagen, H.C., and Cadigan, K.M. (2014). Distinct DNA Binding Sites Contribute to the TCF Transcriptional Switch in C. elegans and Drosophila. PLOS Genetics 10, e1004133.

Bialecka, M., Young, T., Sousa Lopes, S.C. de, ten Berge, D., Sanders, A., Beck, F., and Deschamps, J. (2012). Cdx2 contributes to the expansion of the early primordial germ cell population in the mouse. Developmental Biology 371, 227–234.

Cadigan, K.M. (2012). Chapter one - TCFs and Wnt/β-catenin Signaling: More than One Way to Throw the Switch. In Current Topics in Developmental Biology, S.P. and F. Payre, ed. (Academic Press), pp. 1–34.

Cadigan, K.M., and Waterman, M.L. (2012). TCF/LEFs and Wnt Signaling in the Nucleus. Cold Spring Harb Perspect Biol 4, a007906.

Chang, J.L., Chang, M.V., Barolo, S., and Cadigan, K.M. (2008). Regulation of the feedback antagonist naked cuticle by Wingless signaling. Dev Biol 321, 446–454.

Chen, L., and Capra, J.A. (2020). Learning and interpreting the gene regulatory grammar in a deep learning framework. PLOS Computational Biology 16, e1008334.

Clevers, H., and Nusse, R. (2012). Wnt/β-Catenin Signaling and Disease. Cell 149, 1192–1205.

Clevers, H., Loh, K.M., and Nusse, R. (2014). An integral program for tissue renewal and regeneration: Wnt signaling and stem cell control. Science 346, 1248012.

Concordet, J.-P., and Haeussler, M. (2018). CRISPOR: intuitive guide selection for CRISPR/Cas9 genome editing experiments and screens. Nucleic Acids Res 46, W242–W245.

Doumpas, N., Lampart, F., Robinson, M.D., Lentini, A., Nestor, C.E., Cantù, C., and Basler, K. (2018). TCF/LEF dependent and independent transcriptional regulation of Wnt/β‐catenin target genes. The EMBO Journal e98873.

van Es, J.H., Haegebarth, A., Kujala, P., Itzkovitz, S., Koo, B.-K., Boj, S.F., Korving, J., van den Born, M., van Oudenaarden, A., Robine, S., et al. (2012). A Critical Role for the Wnt Effector Tcf4 in Adult Intestinal Homeostatic Self-Renewal. Mol Cell Biol 32, 1918–1927.

Farley, E.K., Olson, K.M., Zhang, W., Rokhsar, D.S., and Levine, M.S. (2016). Syntax compensates for poor binding sites to encode tissue specificity of developmental enhancers. Proc. Natl. Acad. Sci. U.S.A. 113, 6508–6513.

Fornes, O., Castro-Mondragon, J.A., Khan, A., van der Lee, R., Zhang, X., Richmond, P.A., Modi, B.P., Correard, S., Gheorghe, M., Baranašić, D., et al. (2020). JASPAR 2020: update of the open-access database of transcription factor binding profiles. Nucleic Acids Res 48, D87–D92.

Franz, A., Shlyueva, D., Brunner, E., Stark, A., and Basler, K. (2017). Probing the canonicity of the Wnt/Wingless signaling pathway. PLOS Genetics 13, e1006700.

Fulco, C.P., Munschauer, M., Anyoha, R., Munson, G., Grossman, S.R., Perez, E.M., Kane, M., Cleary, B., Lander, E.S., and Engreitz, J.M. (2016). Systematic mapping of functional enhancer–promoter connections with CRISPR interference. Science 354, 769–773.

Galceran, J., Fariñas, I., Depew, M.J., Clevers, H., and Grosschedl, R. (1999). Wnt3a-/--like phenotype and limb deficiency in Lef1(-/-)Tcf1(-/-) mice. Genes Dev. 13, 709–717.

Giese, K., Amsterdam, A., and Grosschedl, R. (1991). DNA-binding properties of the HMG domain of the lymphoid-specific transcriptional regulator LEF-1. Genes Dev. 5, 2567–2578.

Grant, C.E., Bailey, T.L., and Noble, W.S. (2011). FIMO: scanning for occurrences of a given motif. Bioinformatics 27, 1017–1018.

Harper, S., and Speicher, D.W. (2011). Purification of proteins fused to glutathione S-tranferase. Methods Mol Biol 681, 259–280.

Hatzis, P., Flier, L.G. van der, Driel, M.A. van, Guryev, V., Nielsen, F., Denissov, S., Nijman, I.J., Koster, J., Santo, E.E., Welboren, W., et al. (2008). Genome-Wide Pattern of TCF7L2/TCF4 Chromatin Occupancy in Colorectal Cancer Cells. Mol. Cell. Biol. 28, 2732–2744.

Hoverter, N.P., Ting, J.-H., Sundaresh, S., Baldi, P., and Waterman, M.L. (2012). A WNT/p21 Circuit Directed by the C-Clamp, a Sequence-Specific DNA Binding Domain in TCFs. Mol. Cell. Biol. 32, 3648–3662.

Hoverter, N.P., Zeller, M.D., McQuade, M.M., Garibaldi, A., Busch, A., Selwan, E.M., Hertel, K.J., Baldi, P., and Waterman, M.L. (2014). The TCF C-clamp DNA binding domain expands the Wnt transcriptome via alternative target recognition. Nucleic Acids Res 42, 13615–13632.

Jho, E., Zhang, T., Domon, C., Joo, C.-K., Freund, J.-N., and Costantini, F. (2002). Wnt/β-Catenin/Tcf Signaling Induces the Transcription of Axin2, a Negative Regulator of the Signaling Pathway. Mol. Cell. Biol. 22, 1172–1183.

Jiao, S., Li, C., Hao, Q., Miao, H., Zhang, L., Li, L., and Zhou, Z. (2017). VGLL4 targets a TCF4–TEAD4 complex to coregulate Wnt and Hippo signalling in colorectal cancer. Nature Communications 8, 14058.

Jolma, A., Yin, Y., Nitta, K.R., Dave, K., Popov, A., Taipale, M., Enge, M., Kivioja, T., Morgunova, E., and Taipale, J. (2015). DNA-dependent formation of transcription factor pairs alters their binding specificity. Nature 527, 384–388.

Junion, G., Spivakov, M., Girardot, C., Braun, M., Gustafson, E.H., Birney, E., and Furlong, E.E.M. (2012). A Transcription Factor Collective Defines Cardiac Cell Fate and Reflects Lineage History. Cell 148, 473–486.

Kennedy, M.W., Chalamalasetty, R.B., Thomas, S., Garriock, R.J., Jailwala, P., and Yamaguchi, T.P. (2016). Sp5 and Sp8 recruit β-catenin and Tcf1-Lef1 to select enhancers to activate Wnt target gene transcription. PNAS 113, 3545–3550.

Kent, W.J., Sugnet, C.W., Furey, T.S., Roskin, K.M., Pringle, T.H., Zahler, A.M., and Haussler, and D. (2002). The Human Genome Browser at UCSC. Genome Res. 12, 996–1006.

Kim, K., Cho, J., Hilzinger, T.S., Nunns, H., Liu, A., Ryba, B.E., and Goentoro, L. (2017). Two-Element Transcriptional Regulation in the Canonical Wnt Pathway. Current Biology 27, 2357–2364.e5.

Korinek, V., Barker, N., Morin, P.J., van Wichen, D., de Weger, R., Kinzler, K.W., Vogelstein, B., and Clevers, H. (1997). Constitutive transcriptional activation by a beta-catenin-Tcf complex in APC-/-colon carcinoma. Science 275, 1784–1787.

Kumar, N., Tsai, Y.-H., Chen, L., Zhou, A., Banerjee, K.K., Saxena, M., Huang, S., Toke, N.H., Xing, J., Shivdasani, R.A., et al. (2019). The lineage-specific transcription factor CDX2 navigates dynamic chromatin to control distinct stages of intestine development. Development 146.

Lambert, S.A., Jolma, A., Campitelli, L.F., Das, P.K., Yin, Y., Albu, M., Chen, X., Taipale, J., Hughes, T.R., and Weirauch, M.T. (2018). The Human Transcription Factors. Cell 172, 650–665.

Lengerke, C., Schmitt, S., Bowman, T.V., Jang, I.H., Maouche-Chretien, L., McKinney-Freeman, S., Davidson, A.J., Hammerschmidt, M., Rentzsch, F., Green, J.B.A., et al. (2008). BMP and Wnt Specify Hematopoietic Fate by Activation of the Cdx-Hox Pathway. Cell Stem Cell 2, 72–82.

Leung, J.Y., Kolligs, F.T., Wu, R., Zhai, Y., Kuick, R., Hanash, S., Cho, K.R., and Fearon, E.R. (2002). Activation of AXIN2 Expression by β-Catenin-T Cell Factor. A feedback repressor pathway regulating Wnt signaling. J. Biol. Chem. 277, 21657–21665.

Lewis, A., Freeman-Mills, L., de la Calle-Mustienes, E., Giráldez-Pérez, R.M., Davis, H., Jaeger, E., Becker, M., Hubner, N.C., Nguyen, L.N., Zeron-Medina, J., et al. (2014). A Polymorphic Enhancer near GREM1 Influences Bowel Cancer Risk through Differential CDX2 and TCF7L2 Binding. Cell Reports 8, 983–990.

Lien, W.-H., and Fuchs, E. (2014). Wnt some lose some: transcriptional governance of stem cells by Wnt/β-catenin signaling. Genes Dev. 28, 1517–1532.

Lim, X., Tan, S.H., Yu, K.L., Lim, S.B.H., and Nusse, R. (2016). Axin2 marks quiescent hair follicle bulge stem cells that are maintained by autocrine Wnt/β-catenin signaling. PNAS 113, E1498–E1505.

Lohnes, D. (2003). The Cdx1 homeodomain protein: an integrator of posterior signaling in the mouse. BioEssays 25, 971–980.

Mokry, M., Hatzis, P., Bruijn, E. de, Koster, J., Versteeg, R., Schuijers, J., Wetering, M. van de, Guryev, V., Clevers, H., and Cuppen, E. (2010). Efficient Double Fragmentation ChIP-seq Provides Nucleotide Resolution Protein-DNA Binding Profiles. PLOS ONE 5, e15092.

Moreira, S., Polena, E., Gordon, V., Abdulla, S., Mahendram, S., Cao, J., Blais, A., Wood, G.A., Dvorkin-Gheva, A., and Doble, B.W. (2017). A Single TCF Transcription Factor, Regardless of Its Activation Capacity, Is Sufficient for Effective Trilineage Differentiation of ESCs. Cell Reports 20, 2424–2438.

Nakamura, Y., and Hoppler, S. (2017). Genome-wide analysis of canonical Wnt target gene regulation in Xenopus tropicalis challenges β-catenin paradigm. Genesis 55, n/a-n/a.

Nakamura, Y., Alves, E. de P., Veenstra, G.J.C., and Hoppler, S. (2016). Tissue- and stage-specific Wnt target gene expression is controlled subsequent to β-catenin recruitment to cis-regulatory modules. Development 143, 1914–1925.

Nallamsetty, S., and Waugh, D.S. (2007). A generic protocol for the expression and purification of recombinant proteins in Escherichia coli using a combinatorial His 6 - maltose binding protein fusion tag. Nature Protocols 2, 383–391.

Panne, D., Maniatis, T., and Harrison, S.C. (2007). An Atomic Model of the Interferon-β Enhanceosome. Cell 129, 1111–1123.

Pfaffl, M.W. (2001). A new mathematical model for relative quantification in real-time RT– PCR. Nucleic Acids Res 29, e45.

Pilon, N., Oh, K., Sylvestre, J.-R., Bouchard, N., Savory, J., and Lohnes, D. (2006). Cdx4 is a direct target of the canonical Wnt pathway. Developmental Biology 289, 55–63.

Pilon, N., Oh, K., Sylvestre, J.-R., Savory, J.G.A., and Lohnes, D. (2007). Wnt signaling is a key mediator of Cdx1 expression in vivo. Development 134, 2315–2323.

Pomerantz, M.M., Ahmadiyeh, N., Jia, L., Herman, P., Verzi, M.P., Doddapaneni, H., Beckwith, C.A., Chan, J.A., Hills, A., Davis, M., et al. (2009). The 8q24 cancer risk variant rs6983267 shows long-range interaction with MYC in colorectal cancer. Nat Genet 41, 882–884.

Quinlan, A.R., and Hall, I.M. (2010). BEDTools: a flexible suite of utilities for comparing genomic features. Bioinformatics 26, 841–842.

Ramakrishnan, A.-B., and Cadigan, K.M. (2017). Wnt target genes and where to find them. F1000Res 6, 746.

Ran, F.A., Hsu, P.D., Wright, J., Agarwala, V., Scott, D.A., and Zhang, F. (2013). Genome engineering using the CRISPR-Cas9 system. Nat. Protocols 8, 2281–2308.

Ravindranath, A., and Cadigan, K.M. (2014). Structure-Function Analysis of the C-clamp of TCF/Pangolin in Wnt/ß-catenin Signaling. PLoS One 9.

Savic, D., Partridge, E.C., Newberry, K.M., Smith, S.B., Meadows, S.K., Roberts, B.S., Mackiewicz, M., Mendenhall, E.M., and Myers, R.M. (2015). CETCh-seq: CRISPR epitope tagging ChIP-seq of DNA-binding proteins. Genome Res. 25, 1581–1589.

Schweizer, L., Nellen, D., and Basler, K. (2003). Requirement for Pangolin/dTCF in Drosophila Wingless signaling. Proc. Natl. Acad. Sci. U.S.A. 100, 5846–5851.

Shimizu, T., Bae, Y.-K., Muraoka, O., and Hibi, M. (2005). Interaction of Wnt and caudal-related genes in zebrafish posterior body formation. Developmental Biology 279, 125–141.

Smith, R.P., Taher, L., Patwardhan, R.P., Kim, M.J., Inoue, F., Shendure, J., Ovcharenko, I., and Ahituv, N. (2013). Massively parallel decoding of mammalian regulatory sequences supports a flexible organizational model. Nat Genet 45, 1021–1028.

Söderholm, S., and Cantù, C. (2021). The WNT/β-catenin dependent transcription: A tissue-specific business. WIREs Systems Biology and Medicine *n/a*, e1511.

Spitz, F., and Furlong, E.E.M. (2012). Transcription factors: from enhancer binding to developmental control. Nat Rev Genet 13, 613–626.

Stamos, J.L., and Weis, W.I. (2013). The β-Catenin Destruction Complex. Cold Spring Harb Perspect Biol 5, a007898.

Swanson, C.I., Evans, N.C., and Barolo, S. (2010). Structural Rules and Complex Regulatory Circuitry Constrain Expression of a Notch- and EGFR-Regulated Eye Enhancer. Developmental Cell 18, 359–370.

Thanos, D., and Maniatis, T. (1995). Virus induction of human IFNβ gene expression requires the assembly of an enhanceosome. Cell 83, 1091–1100.

Tomlinson, I., Webb, E., Carvajal-Carmona, L., Broderick, P., Kemp, Z., Spain, S., Penegar, S., Chandler, I., Gorman, M., Wood, W., et al. (2007). A genome-wide association scan of tag SNPs identifies a susceptibility variant for colorectal cancer at 8q24.21. Nature Genetics 39, 984–988.

Tuupanen, S., Turunen, M., Lehtonen, R., Hallikas, O., Vanharanta, S., Kivioja, T., Björklund, M., Wei, G., Yan, J., Niittymäki, I., et al. (2009). The common colorectal cancer predisposition SNP rs6983267 at chromosome 8q24 confers potential to enhanced Wnt signaling. Nat Genet 41, 885–890.

Valenta, T., Hausmann, G., and Basler, K. (2012). The many faces and functions of β‐catenin. The EMBO Journal 31, 2714–2736.

Van der Flier, L.G., Sabates–Bellver, J., Oving, I., Haegebarth, A., De Palo, M., Anti, M., Van Gijn, M.E., Suijkerbuijk, S., Van de Wetering, M., Marra, G., et al. (2007). The Intestinal Wnt/TCF Signature. Gastroenterology 132, 628–632.

van Amerongen, R., Bowman, A.N., and Nusse, R. (2012). Developmental Stage and Time Dictate the Fate of Wnt/β-Catenin-Responsive Stem Cells in the Mammary Gland. Cell Stem Cell 11, 387–400.

Verzi, M.P., Hatzis, P., Sulahian, R., Philips, J., Schuijers, J., Shin, H., Freed, E., Lynch, J.P., Dang, D.T., Brown, M., et al. (2010). TCF4 and CDX2, major transcription factors for intestinal function, converge on the same cis-regulatory regions. PNAS 107, 15157–15162.

Verzi, M.P., Shin, H., Roman, A.K.S., Liu, X.S., and Shivdasani, R.A. (2013). Intestinal Master Transcription Factor CDX2 Controls Chromatin Access for Partner Transcription Factor Binding. Molecular and Cellular Biology 33, 281–292.

Wang, B., Zhao, L., Fish, M., Logan, C.Y., and Nusse, R. (2015). Self-renewing diploid Axin2+ cells fuel homeostatic renewal of the liver. Nature 524, 180–185.

Wasserman, W.W., and Sandelin, A. (2004). Applied bioinformatics for the identification of regulatory elements. Nature Reviews Genetics 5, 276–287.

van de Wetering, M., Sancho, E., Verweij, C., de Lau, W., Oving, I., Hurlstone, A., van der Horn, K., Batlle, E., Coudreuse, D., Haramis, A.-P., et al. (2002). The β-Catenin/TCF-4 Complex Imposes a Crypt Progenitor Phenotype on Colorectal Cancer Cells. Cell 111, 241–250.

Wright, J.B., Brown, S.J., and Cole, M.D. (2010). Upregulation of c-MYC in cis through a Large Chromatin Loop Linked to a Cancer Risk-Associated Single-Nucleotide Polymorphism in Colorectal Cancer Cells. Mol. Cell. Biol. 30, 1411–1420.

Yochum, G.S., Cleland, R., and Goodman, R.H. (2008). A Genome-Wide Screen for β-Catenin Binding Sites Identifies a Downstream Enhancer Element That Controls c-Myc Gene Expression. Molecular and Cellular Biology 28, 7368–7379.

Young, T., Rowland, J.E., van de Ven, C., Bialecka, M., Novoa, A., Carapuco, M., van Nes, J., de Graaff, W., Duluc, I., Freund, J.-N., et al. (2009). Cdx and Hox Genes Differentially Regulate Posterior Axial Growth in Mammalian Embryos. Developmental Cell 17, 516–526.

Zanke, B.W., Greenwood, C.M., Rangrej, J., Kustra, R., Tenesa, A., Farrington, S.M., Prendergast, J., Olschwang, S., Chiang, T., Crowdy, E., et al. (2007). Genome-wide association scan identifies a colorectal cancer susceptibility locus on chromosome 8q24. Nature Genetics 39, 989–994.

Zimmerli, D., Borrelli, C., Jauregi-Miguel, A., Söderholm, S., Brütsch, S., Doumpas, N., Reichmuth, J., Murphy-Seiler, F., Aguet, Mi., Basler, K., et al. (2020). TBX3 acts as tissue-specific component of the Wnt/β-catenin transcriptional complex. ELife 9, e58123.

